# FplA From *Fusobacterium nucleatum* is a Type Vd autotransporter phospholipase with a proposed role in altered host signaling and evasion of autophagy

**DOI:** 10.1101/136598

**Authors:** Michael A. Casasanta, Christopher C. Yoo, Hans B. Smith, A. Jane Duncan, Kyla Cochrane, Ann C. Varano, Emma Allen-Vercoe, Daniel J. Slade

## Abstract

*Fusobacterium nucleatum* is a pathogenic oral bacterium that is linked to multiple human infections and colorectal cancer. While most Gram-negative pathogens utilize secretion systems for cellular invasion and infection, *F. nucleatum* lacks Type I, II, III, IV, and VI secretion. By contrast, *F. nucleatum* strains are enriched in Type V secreted autotransporters, which are Gram-negative bacterial virulence factors critical for binding and entry into host cells. Here we present the first biochemical characterization of a *F. nucleatum* Type Vd phospholipase class A1 autotransporter (strain ATCC 25586, gene FN1704) that we hereby rename *Fusobacterium* phospholipase autotransporter (FplA). FplA is expressed as a full-length 85 kDa outer membrane embedded protein, or as a truncated phospholipase domain that remains associated with the outer membrane. Using multiple FplA constructs we characterized lipid substrate specificity, potent inhibitors, and chemical probes to detect and track this enzyme family. While the role of FplA is undetermined in *F. nucleatum* virulence, homologous phospholipases from intracellular pathogens are critical for vacuole escape, altered host signaling, and intracellular survival. We hypothesize that upon intracellular invasion of the host, FplA could play a role in phagosomal escape, subversion of autophagy, or eicosanoid-mediated inflammatory signaling, as we show that FplA binds with high affinity to host phosphoinositide signaling lipids critical to these processes. Our identification of substrates, inhibitors, and chemical probes for FplA, in combination with an *fplA* gene deletion strain, encompass a powerful set of tools for the future analysis of FplA *in vivo*. In addition, these studies will guide the biochemical characterization of additional Type Vd autotransporter phospholipases.

**IMPORTANCE:** *F. nucleatum* is an emerging pathogen that is linked to the pathogenesis of colorectal cancer, yet there is a critical knowledge gap in the mechanisms used by this bacterium to elicit changes in the host for intracellular entry and survival. As phospholipases are critical virulence factors for intracellular bacteria to initiate vacuole lysis, cell-to-cell spread, and evasion of autophagy, we set out to characterize a unique Type Vd secreted phospholipase A1 enzyme from *F. nucleatum*. Our results show a potential role for modulating host signaling pathways through cleavage of phosphoinositide dependent signaling lipids. These studies open the door for further characterization of this unique enzyme family in bacterial virulence, host-pathogen interactions, and for *F. nucleatum*, in colorectal carcinogenesis.

## INTRODUCTION

*Fusobacterium nucleatum* is an emerging oral pathogen that readily disseminates, presumably through hematogenous spread^1,2^, to cause potentially fatal infections of the brain^3^, liver^4^, lungs^5^, heart^6^, appendix^7^, and amniotic fluid where it causes pre-term birth^1,8,9^. Recent studies have uncovered a correlation between colorectal cancer tumors and an overabundance of *F. nucleatum* present in diseased tissue^10–12^. Subsequent studies confirmed a potential causative effect for *F. nucleatum* in the onset and progression of disease using an APC ^min/-^ mouse model of accelerated CRC pathogenesis, where upon oral gavage with *F. nucleatum,* mice showed increased numbers of intestinal tumors^13^. Subsequent experiments show that intravenous injection of *F. nucleatum* results in bacterial localization to mouse tumor tissues rich in Gal-GalNAc surface polysaccharide in a Fap2 autotransporter protein dependent manner^2^. In addition, human patients that had the highest detected levels of *F.nucleatum* within tumors had the lowest survival rate^14^. Invasive *F. nucleatum* strains can enter into epithelial and endothelial cells^15,16^, which induces the secretion of proinflammatory cytokines that drive local inflammation as seen in colorectal cancer^13^, and also provides a niche in which the bacterium can subvert the host immune system. Previously characterized proteins involved in host cell binding and invasion include FadA (ATCC 25586, gene FN0264), a small helical adhesin that binds to E-cadherin and modulates prevalent colorectal cancer signaling pathways^17,18^; Fap2 (ATCC 25586, gene FN1449), a galactose inhibitable Type Va secreted adhesin that binds Gal-GalNAc sugars^2,19–21^; and RadD (ATCC 25586, gene 1526), an arginine inhibitable Type Va autotransporter adhesin^19,22^. Upon interaction with oral epithelial cells, *F. nucleatum* also induces the production of human β-defensin 2 and 3 (hBD2, hBD3), which are secreted, cationic antimicrobial peptides that can directly kill Gram-negative bacteria, and also act as chemo-attractants to modulate adaptive immunity during infection^23,24^. Despite our knowledge of the intracellular and immune modulating lifestyle of *F. nucleatum*, very few proteins have been characterized that play a role in intracellular survival and subversion of bacterial clearance systems such as autophagy.

*F. nucleatum* is unique among pathogenic bacteria in that it does not harbor large, multi-protein secretion systems (Types I-IV, VI, and IX in Gram-negative bacteria) to establish infections and alter host-signaling for survival^25^. To compensate for this apparent lack of virulence factors, invasive and isolated clinical strains of *F. nucleatum* contain an overabundance of uncharacterized proteins containing type II membrane occupation and recognition nexus (MORN2) domains, and a genomic expansion of Type V secreted effectors known as autotransporters^16^. Autotransporters are large outer membrane and secreted proteins that are divided into 5 classes (Type Va-Ve) based on their domain architecture, and are critical proteins in host cell adherence, invasion, and biofilm formation^26,27^. The type Vd autotransporter PlpD from *Pseudomonas aeruginosa* was recently characterized biochemically and structurally and revealed a secreted N-terminal patatin-like protein (PFAM: PF01734) with an α-β hydrolase fold containing a catalytic dyad (Ser, Asp) conferring phospholipase A1 activity (EC 3.1.1.32) through the hydrolysis of glycerophospholipid moieties at the *sn*-1 position to release a fatty acid^28,29^. In addition, PlpD contains a 16-strand C-terminal Beta barrel domain of the bacterial surface antigen family (PFAM: PF01103) for initial outer membrane anchorage, and a predicted periplasmic polypeptide-transport-associated (POTRA) domain potentially involved in protein folding and translocation of the phospholipase domain to the surface. The PlpD secreted phospholipase domain was able to disrupt liposomes and was also shown to bind multiple phospholipids, including the phosphoinositide class of human intracellular signaling lipids^30^. Upon our analysis, we found that in most cases, *F. nucleatum* genomes each contain one gene (in strain ATCC 25586, gene FN1704, UniProtKB-Q8R6F6 - herein renamed *fplA*) encoding for a previously uncharacterized type Vd autotransporter that is homologous to PlpD.

Bacterial phospholipases play critical roles in virulence by converting vacuoles or phagosomes into protective encasements for replication and survival, or by aiding in vacuole lysis to achieve liberation into the cytoplasm and subversion of host lysosomal induced death^31,32^. Bacterial acyl hydrolases of the A, B, and lysophospholipase (LPAs) families are almost exclusively membrane-associated or injected into the host by secretion systems, many through the well-characterized type 3 secretion system (T3SS)^33^. Intracellular pathogens including *Helicobacter*, *Listeria*, *Salmonella*, *Shigella*, *Pseudomonas, and Legionella* rely on phospholipases for intracellular survival and intercellular spread^32^. *Helicobacter pylori* uses the outer membrane phospholipase PldA1 for virulence, and this enzyme is involved in growth at low pH^34^, colonization of the gastric mucosa^35^, and hemolytic activity^35^. *Listeria monocytogenes* secretes two phospholipase C proteins (PI-PLC and PC-PLC), and these enzymes are critical for late time point evasion of autophagy and establishment of an intracellular lifestyle^36^. Structural predictions of FplA align well with the *Pseudomonas aeruginosa* type III secreted toxin ExoU, which is activated by ubiquitination and also binds to ubiquitinated host proteins to initiate its phospholipase A2 and lysphospholipase activity_37_, thereby causing PI(4,5)_2_-associated cytoskeletal collapse and arachadonic acid dependent inflammatory signaling^38,39^. In addition, FplA is highly homologous to the *Legionella pneumophila* effector VipD, which is a phospholipase A1 enzyme that is activated after binding the human GTPase Rab5^40^ or Rab22, whereupon it protects the bacteria from endosomal fusion and phagosomal maturation in macrophages^41^, and also blocks host apoptosis by cleaving mitochondrial phospholipids^42^. It is noteworthy that there is no evidence that *F. nucleatum* induces cytoskeletal collapse or cell death upon infection of epithelial or endothelial cells; this has led us to hypothesize a role for FplA more similar to VipD in evasion of autophagy.

Understanding the molecular mechanisms used by *F. nucleatum* to divert intracellular clearance will provide tools to dissect host-pathogen interactions critical for persistent infection and modulation of cell-signaling pathways as seen associated with colorectal cancer. While multiple mechanisms of cellular binding and entry have been identified, there are no studies to determine if this initial binding needs to be aided by additional enzymes or factors for breaching the host membranous barrier. We propose a model in which FplA has the enzymatic potential to perform a diverse set of functions in *F. nucleatum* virulence and intracellular colonization, including lipid cleavage for entry into host cells, vacuole escape for cytoplasmic access, and cleavage of phosphoinositide signaling lipids to subvert host defense mechanisms including autophagy.

## RESULTS

### FN1704 encodes for a type Vd phospholipase autotransporter

*Fusobacterium* phospholipase autotransporter (FplA, UniProtKB-Q8R6F6) was identified as the gene previous labeled FN1704 in *F. nucleatum* ATCC 25586. Domain identification was carried out using SignalP 4.1 to identify a signal sequence (residues 1-19), and the SWISS-MODEL^43^ structure prediction server which allowed the identification of a patatin domain responsible for phospholipase activity (residues 60-350), a POTRA domain common in protein-protein interactions (residues 351-431), and a C-terminal beta barrel domain (residues 431-760) to insert FplA in the outer membrane. In addition, we identified a unique 40 amino acid N-terminal extension (NTE, residues 20-59) that plays a role in the catalytic efficiency of the enzyme likely by being critical for proper protein folding and position of the active site residues, and not substrate binding (**Fig. 1A, Fig. S1A**). Structure prediction of this enzyme revealed the N-terminal patatin domain is highly similar to PlpD from *Pseudomonas aeruginosa* (PDB: 5FQU), and an alignment shows an overall fold in residues 60-343 (32% identity corresponding to PlpD residues 22-311) that upon alignment globally have a root mean squared deviation (RMSD) of 0.20 angstroms, with a highly conserved active site containing a catalytic dyad (Ser98 and Aps243) and an oxyanion hole (Gly69/70/71) (**Fig. 1B, Fig. S1B**). In addition, the next closest structural homologs of the FplA catalytic domain (residues 60-343) are predicted to be the non-autotransporter phospholipase A enzymes ExoU (Type III secreted) from *P. aeruginosa* (19.0% identity to residues 102-472, PDB: 4AKX, 3TU3) and VipD (Type IV secreted) from *Legionella pneumophila* (17.3% identity to residues 33-411, PDB: 4AKF), with an overall RMSD of 10.8 Å for ExoU and 5.0 Å for VipD for structural alignments (**Fig. S2A-B**).

**Fig 1.**
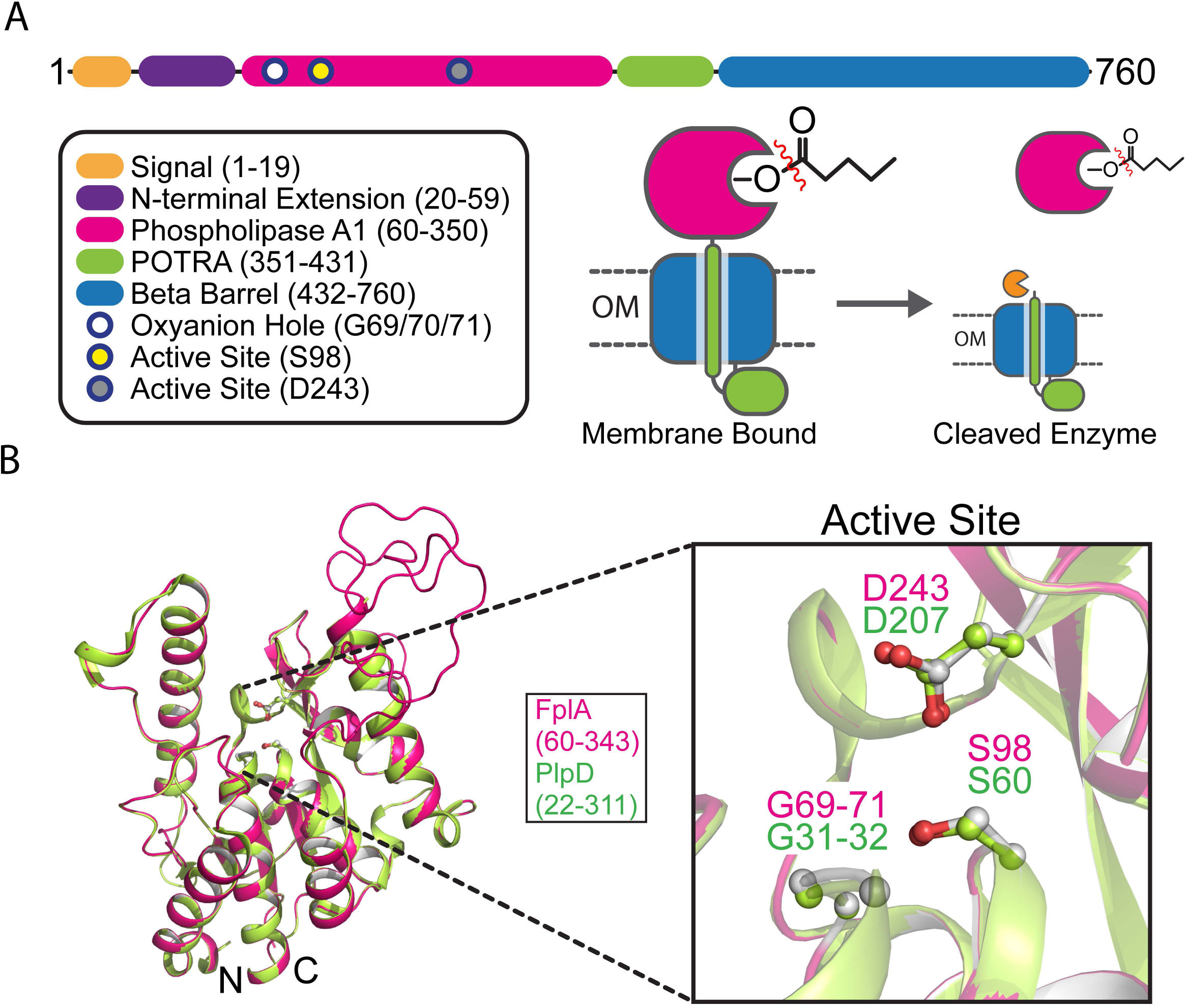
FplA is a Type Vd autotransporter phospholipase from *Fusobacteirum nucleatum.* (A) Structure and description of FplA domains and their location in the periplasm, outer membrane, and surface exposure of the phospholipase A1 (PLA_1_) domain. (B) Alignment of a predicted FplA PLA_1_ domain structure overlayed with the crystal structure (PDB: SFQU) of the homogous enzyme PlpD from Pseudomonas Aeruginosa, with a magnified view of the catalytic dyad (S98, D243) and oxyanion hole (G69, G70, G71).

### Characterization of fluorogenic substrates to probe the phospholipase A1 (PLA_1_) activity of FplA

Multiple FplA constructs were cloned from the *F. nucleatum* 25586 genome and expressed in *E. coli*, including variations that lack a signal sequence for cytoplasmic expression (Residues 20-431, 20-350, 60-431, 60-350), and a full-length version in which we replaced the native signal sequence with an *E. coli* OmpA signal for more robust expression and surface presentation (OmpA 1-27/FplA 20-760). Constructs were tested for their phospholipase activity using substrates specific for either A1 or A2 class enzymes, as the homolog PlpD from *P. aeruginosa* showed specific A1 activity. We showed that FplA has only PLA_1_ activity (**Fig. 2A**, **Fig. S3A-B**) using the PLA_1_ specific substrate PED-A1, and further demonstrated that the general lipase substrates 4-Methyl Umbelliferyl Butyrate (4-MuB) and 4-Methyl Umbelliferyl Heptanoate (4-MuH) are robust tools for studies of FplA and other Type Vd autotransporter phospholipases (**Fig. 2A**). In addition we determined this enzyme is not dependent on calcium for activity (**Fig. S3C**), and that it is most active at pH 8.5 (**Fig. S3D**). The first full Michaelis-Menten kinetics for a Type Vd autotransporter were performed on each FplA construct using 4-MuH as a substrate, and indicated that amino acids 20-350, incorporating the N-terminal extension and catalytic PLA_1_ domain, shows the most robust catalytic efficiency (*k*_cat_/*K*_m_ = 3.2 x 10^6^ s^-1^ M^-1^) (**Fig. 2B, Fig. S3E**). Upon removal of the N-terminal extension, constructs had lower substrate turnover rates (*k*_cat_), but the relative binding affinities (*K*_m_) for 4-MuH was unchanged (**Fig. 2C-F**). We also show that tighter binding was seen with the substrate that most closely mimics a phospholipid (PED-A1, *K*_m_=1.90 μM), and of the single acyl chain substrates, 4-MuH (7 carbon acyl chain) resulted in significantly tighter binding (*K*_m_=19 μM) than with the 4 carbon acyl chain substrate 4-MuB (*K*_m_=500 μM) (**Fig. 2D**). Mutation of the active site serine (S98A) and aspartate (D243A) residues that make up the catalytic dyad resulted in no detectable enzymatic activity (**Fig. 2B**). In addition, the glycine rich stretch that constitutes the oxyanion hole (G69/70/71) was analyzed, but G→A single mutations or multiple glycine changes (G69/70/71A) rendered the proteins insoluble (unpublished data) and therefore could not be used for enzymatic analysis.

**Fig 2.**
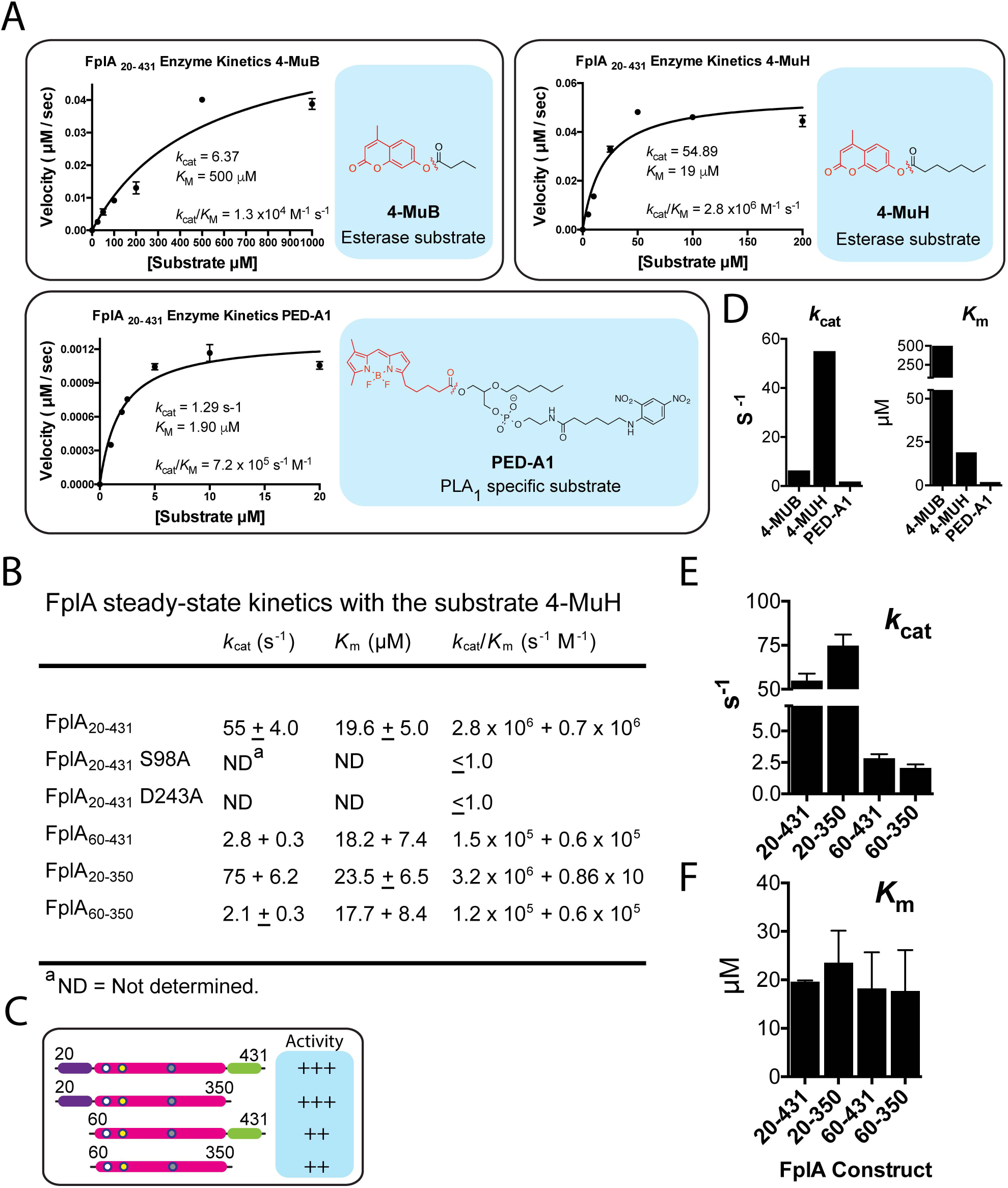
Characterization of FplA lipase activity with multiple fluorescent substrates. (A) FplA is a PLA1 specif ic enzyme as shown by cleavage of the substrate PED-A 1. FplA also efficiently processes the saturated acyl chain of the substrates 4-MuB (4-Methyl Umbelliferyl Butyrate) and 4-MuH (4-Methyl Umbelliferyl Hepta noate). (B) Steady-state kinetics of multiple FplA constructs with 4-MuH. (C) Visual representation of FplA catalytic acivity when expressed as truncated versions lacking specific domains. (D-F) Characterization of FplA turnover rates and substrate binding affinities with multiple substrates. Results show FplA binds longer acyl chains with higher affinity, adn that loss of the N-terminal Extension domain (residues 20-59) reduces turnover rate but does not affect substrate binding affinity.

### Identification of FplA inhibitors and chemical probes for *in vitro* enzyme characterization

We present the first characterization of inhibitors for Type Vd autotransporter phospholipases. Our library was chosen based on previously characterized compounds used to inhibit a broad range of lipases. We show that the classic calcium-dependent PLA_2_ inhibitor Methyl Arachidonyl Fluorophosphonate (MAFP) is the most potent for FplA with an IC_50_ of 11 nM^44^. Additional potent inhibitors contained a trifluoromethyl ketone head-group (ATFMK) which also covalently binds to active site serines within enzymes, or an enylfluorophosphonate group (**Fig. 3A-C**). We observed that IDEFP is a much more potent inhibitor than IDFP; these compounds differ by only a double bond at the end of the IDEFP acyl chain. In addition, MAFP is the most potent inhibitor and the arachidonyl portion of the molecule contains four double bonds, making it and ATFMK the most unsaturated substrates of the inhibitors tested. We therefore hypothesize that FplA binds and docks unsaturated acyl chain substrates and inhibitors with much higher affinity than saturated acyl chains, potentially because of angular changes within the molecule at double bonds. Additional inhibitors were tested that showed no significant activity against FplA (IC_50_ > 100 μM) and their analysis, as well as IC_50_ plots for all inhibitors are presented in **Fig. S4**. Among the non-potent inhibitors are LY311727, an inhibitor of human secretory PLA_2_ (sPLA_2_)_45_, and Manoalide, an inhibitor human group II sPLA_2_, cobra venom PLA2, and phospholipase C (PLC)^46–48^. These inhibitors are predicted to not be competitive inhibitors that irreversibly inactivate the catalytic serines, and therefore are likely specific for motifs not present in FplA.

**Fig 3.**
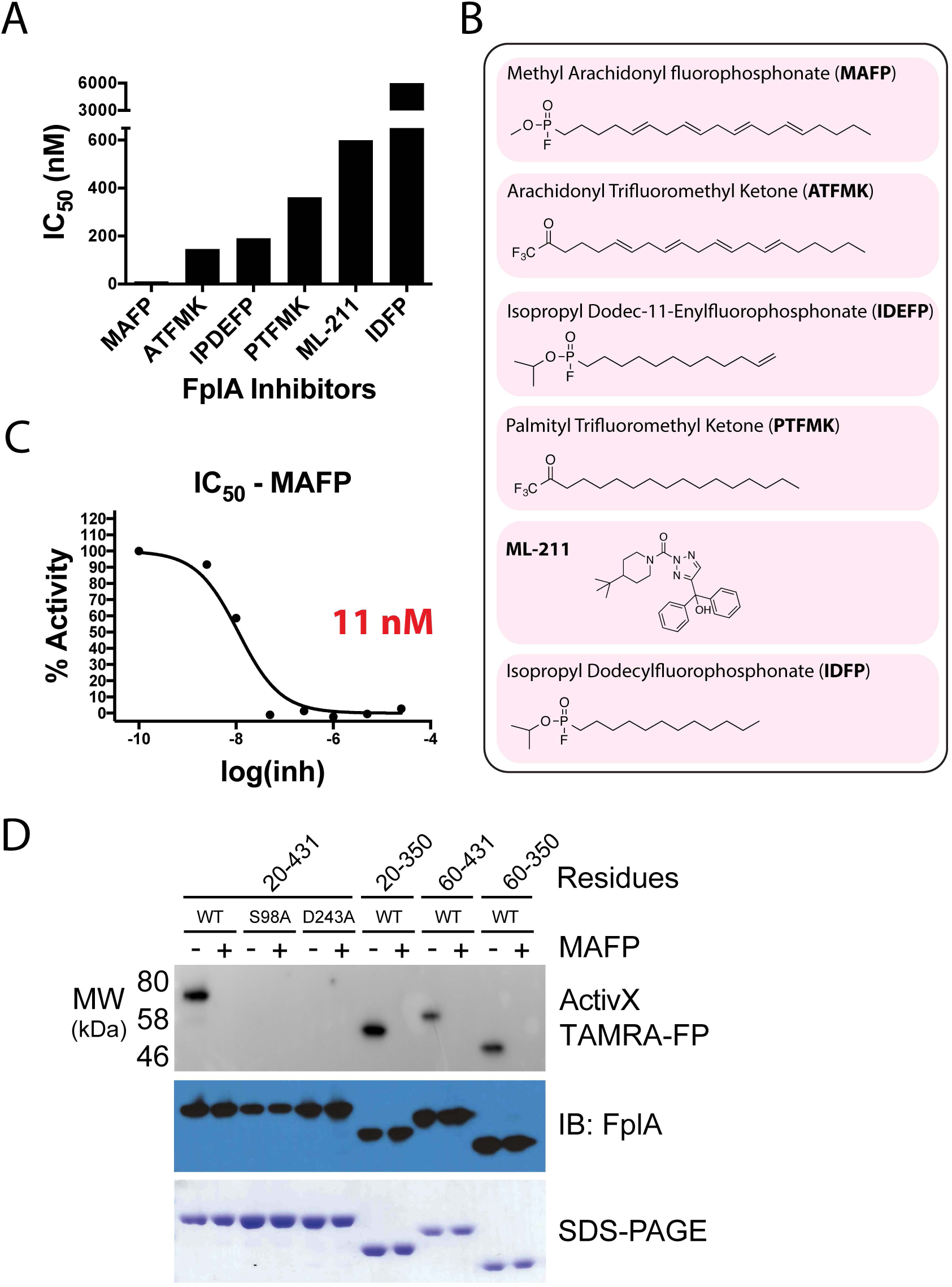
Characterization of FplA inhibitors. (A) IC_50_ assays showing varying degrees of inhibition towards FplA by inhibitors previously shown to inhibit a variety of Iipases. (B) Structure and names of tested inhibi tors. (C) IC_50_ plot of MAFP, the most potent (11 nM) FplA inhibitor charactrerized. (D) Analysis of the active site of FplA shows that the active site serine (S98) reacts with ActivX TAMRA-FP probe, but can not bind in the presence of the competitive inhibitor MAFP. S98A and D243A mutants will not bind the serince active site probe. Western blot and SOS-PAGE gels stained with Coomassie blue serve as load controls for all constructs.

Activity-based protein profiling (ABPP) probes have been used extensively to study the serine hydrolase super-family of proteins^49,50^. The ActivX TAMRA-FP probe, which contains a fluorophosphonate (FP) headgroup to label the active site serine, along with a linker and a fluorescent TAMRA molecule, was used to label purified FplA constructs (**Fig. 3D**). The ActivX TAMRA-FP probe labeled active FplA, but did not bind to the S98A or D243A mutants. We propose that in the absence of D243 which stabilizes substrate, the probe does not properly interact with S98 to initiate covalent labeling. In addition, in the presence of the inhibitor MAFP, the ActivX TAMRA-FP probe is unable to bind to FplA due to competitive inhibition (**Fig. 3D**). We further demonstrated that load controlling is even by transferring the probe bound proteins to PVDF and immunoblotting using a custom FplA antibody that we generated in rabbits using the FplA_20-431_ construct as an antigen.

### Expression of full-length FplA on the surface of *E. coli*

As PLA_1_ enzymes are increasingly being recognized as important virulence factors, the discovery that all bioinformatically identified Type Vd autotransporters belong to this enzyme family is potentially significant^28^. Previously described work on the characterization of PlpD from *P. aeruginosa* uncovered an enzyme that is cleaved and released into the media. We created a FplA construct from *F. nucleatum* 25586 for recombinant expression in *E. coli* that removed the native signal sequence (residues 1-19) and replaced it with the signal sequence from *E. coli* OmpA (residues 1-27). This resulted in more robust expression of FplA on the surface of *E. coli* when compared with using a native signal sequence, which may not be recognized as efficiently by the *E. coli* Sec machinery (native signal sequence data not shown). We demonstrated that FplA can be efficiently exported through the Sec apparatus, assembled in the outer membrane, and the PLA_1_ domain of FplA is present and functional on the surface of *E. coli*. In **Fig. 4A** we show that full-length FplA was detected on the surface of *E. coli* by fluorescence microscopy when using our polyclonal FplA antibody for detection and an Alexa Fluor 488 conjugated secondary antibody. FplA on the surface was active as addition of whole live bacteria to a reaction containing the fluorogenic substrate 4-MuH resulted in cleavage of the lipid substrate and a subsequent increase in fluorescence, which was inhibited by the addition of MAFP (**Fig. 4B**). To further prove that full-length FplA is expressed on the surface of E. coli, we confirm that treatment with the non-specific and cell-impermeable protease, Proteinase K (PK), cleaves FplA from the surface, but does not cleave the cytoplasmic control GAPDH (**Fig. 4C-D**).

**Fig 4.**
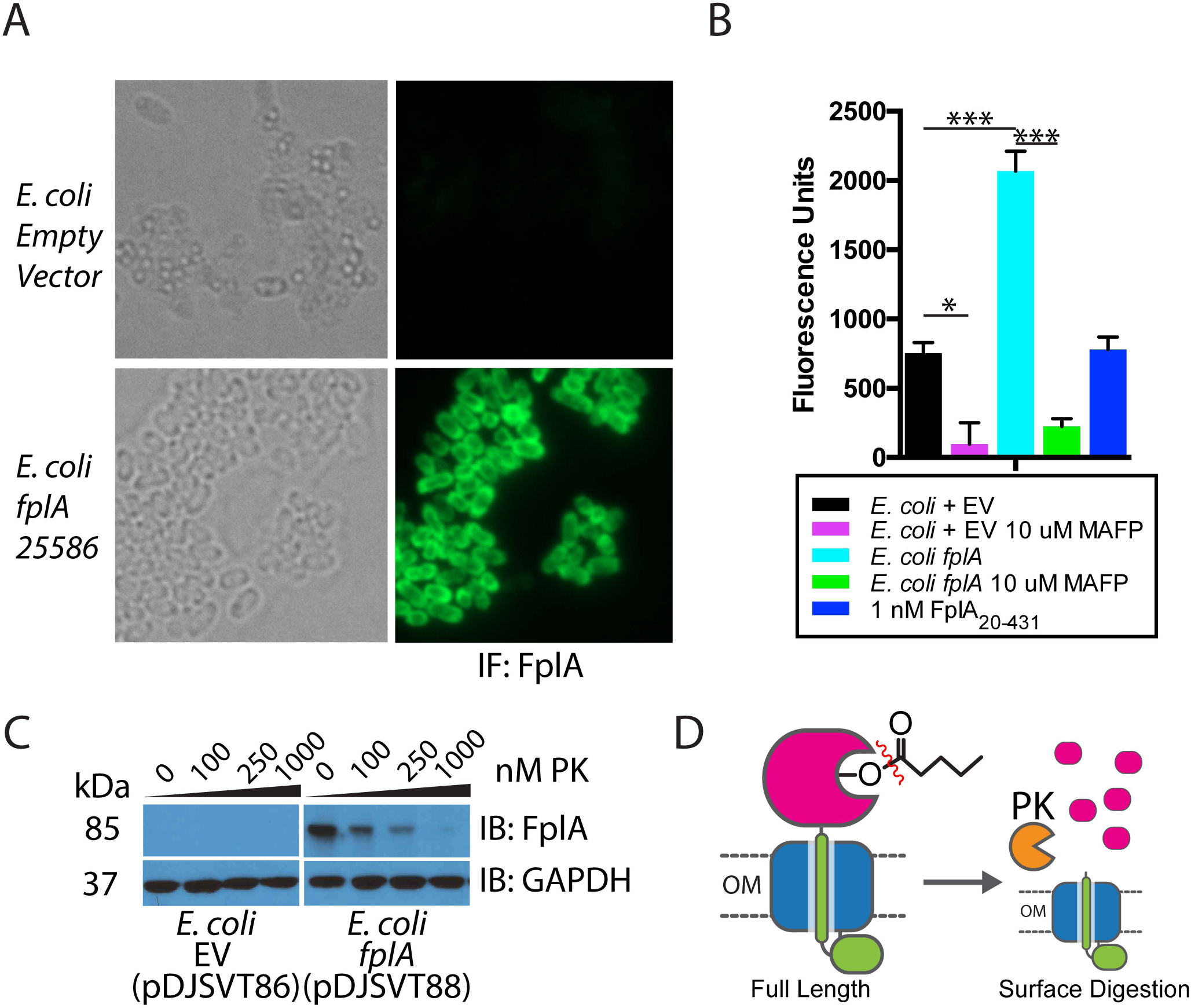
Expression of Full length FplA inf.*coli* and functional analysis. (A) An OmpA_1-27_ signal sequence allows for robust expression of FplA_20-760_ on the surface of *E.coli* as seen by fluorescence microscopy with an anti-FplA antibody. (B) Enzymatic activity of FplA when livef.*coli* are added to reactions containing 4-MuH as a fluorescent substrate (C) Proteinase K (PK), a cell impenetrable non-specific protease, is able to digest surface exposed FplA, but not the cytoplasmic protein GAPGH. (D) Schematic of PK cleavage of full-length FplA from the surface of *E.coli.* EV= Empty vector. Statistical analysis was performed using a multiple comparison analysis by one-way ANOVA. p-values: * = ≤ 0.05, *** = ≤ 0.0005.

Attempts to detect FplA on the surface of *F. nucleatum* 23726 and *F. nucleatum* 25586 by fluorescence microscopy were unsuccessful, which we attribute to the low abundance of this protein as indicated by the need to use large cell quantities in order to see the protein via western blot. It is possible that this is because FplA is such a potent phospholipase that high expression of the enzyme could be detrimental to *F. nucleatum*, as it could result in self-lysis and cell death. Additionally, we were unable to detect enzymatic activity by placing wild-type *F. nucleatum* 23726 directly in a mixture of 4-MuH substrate (results not shown). Neither of these negative results for activity were surprising considering the low amount of FplA present; such a lack of activity at the surface is not uncommon for other outer membrane phospholipases in Gram-negative bacteria. For example, outer membrane phospholipase A (OMPLA) from *E. coli* displays no activity in the absence of outer membrane destabilization compounds such as polymyxin B^51^.

### Creation of an *fplA* deletion strain in *F. nucleatum* 23726

Genetic manipulation of *Fusobacterium* spp. is technically challenging, and of the seven strains used for analysis in this manuscript, only *F. nucleatum* 23726 and 10953 have been successfully mutated by gene deletion^52^. A single homologous crossover plasmid (pDJSVT100, **Table S2**) that we developed from a *Clostridium* shuttle vector^53^ using a recombination method previously established for *F. nucleatum*^*52*^ was used to create a Δ*fplA* strain (Gene HMPREF0397_1968) (Strain DJSVT01, **Table S1**) marked with chloramphenicol resistance (**Fig. 5A-B**). We verified by PCR that the *fplA* gene was disrupted by the chromosomally-inserted plasmid, and further showed expression of the protein had been abolished by a fluorescent probe and western blots probed with an anti-FplA antibody. As phospholipases have been shown to play a role in bacterial membrane maintenance, we tested *F. nucleatum* 23726 Δ*fplA* for changes in growth rates and cell size, and found that when compared to wild-type *F. nucleatum* 23726, there were no changes in these physical parameters when grown under standard laboratory conditions (**Fig. S5A-D**).

**Fig 5.**
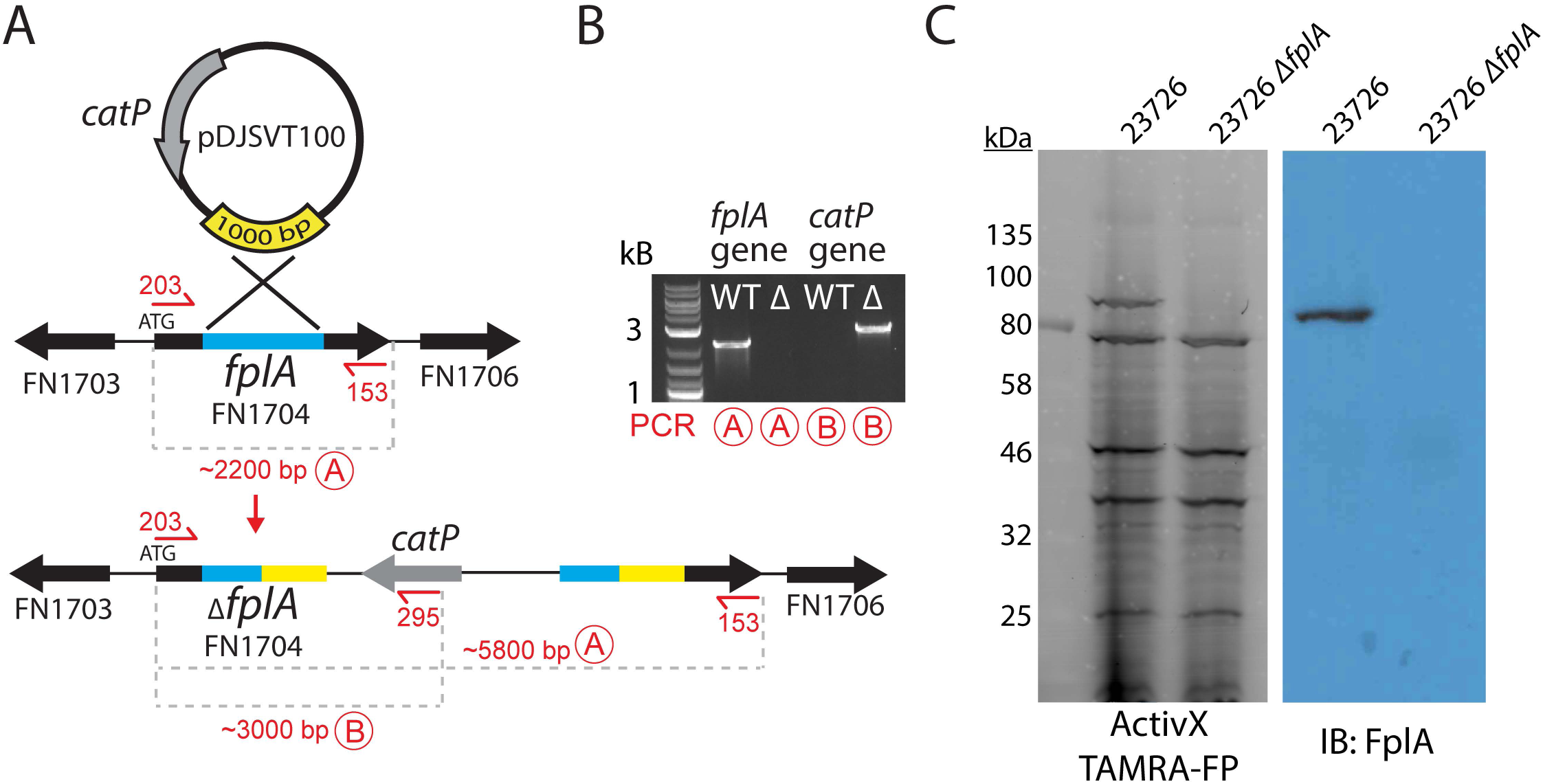
Creation of an *F. nucleatum* 23726 Δ*fplA* strain. (A) pDJSVTl 00 is a single-crossover integration plasmid for disruption of the *fplA* gene. Primers are labeled in red for PCR reactions A and B to confirm plasmid integration and gene knockout. (B) PCR confirmation of the *F. nucleatum* 23726 Δ*fplA* strain. (C) Analysis of FplA protein (85.6 kDa) in WT and Δ*fplA* by fluorescent chemical probe (ActivX TAMRA-FP) to label all active site serine enzymes in the bacteria (Also serves as a load control), followed by transfer to PVDF for western blot analysis by probing with an anti-FplA antibody.

### *F. nucleatum* strains express FplA as a full-length outer membrane protein or as a cleaved phospholipase domain that remains associated with the bacterial surface

Our initial results showed that FplA from *F. nucleatum* 23726 was expressed as a full-length 85 kDa protein, with no apparent release of the PLA_1_ domain from the beta barrel domain. Since PlpD from *P. aeruginosa* is a Type Vd autotransporter that releases the PLA_1_ domain into the media, we sought to see if FplA from seven different *F. nucleatum* strains had different expression patterns or actual physical differences in the size or location of expressed and/or secreted domains. Various FplA proteins were expressed as either a full-length 85 kDa proteins (Strains 23726 and 25586) or as a truncated phospholipase domain (FplA antibody developed against the PLA_1_ and POTRA domains) around 25-30 kDa for strains 10953, 4_8, 4_1_13, 49256, and 7_1 when expressed in either mid-exponential (OD_600_ = 0.7) or stationary phase (OD_600_ = 1.2) (**Fig. 6A**). Interestingly, we could not detect any secreted FplA in the spent culture media, as was previously seen for PlpD from *P. aeruginosa* (**Fig. 6B**). We then tested for the presence of full length FplA in 10953 (cleaved) and 23726 (uncleaved) in early exponential growth (OD_600_ = 0.2) and found that we could detect full length and truncated FplA from 10953, indicating that upon increases in bacterial cell density, FplA is cleaved from the surface by an unknown protein and mechanism (**Fig. 6C**). It is possible that FplA cleavage from the surface results in an active PLA_1_ domain that remains associated with the surface until released by undetermined host factors (pH, molecular cues, etc.) while colonizing specific regions of the human body.

**Fig 6.**
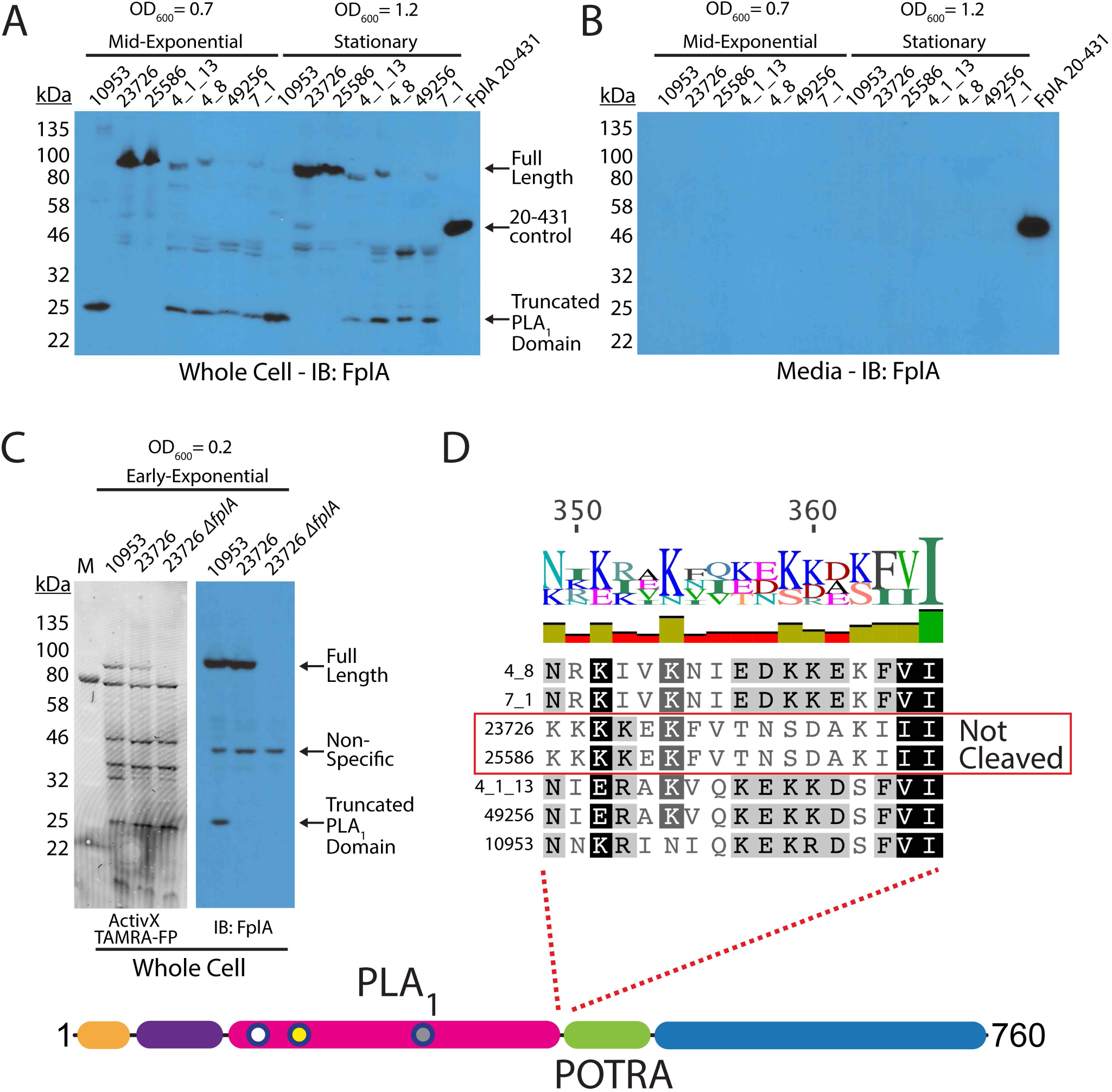
Western blot analysis of FplA in multiple *Fusobacterium* strains. (A) Initial characterization of FplA expression and protein size in mid-exponential phase (OD_600_ = 0.7) and stationary phase (OD_600_ = 1.2) shows that several strains produce a truncated form of FpA that consists of the PLA, domain to which the FplA antibody was raised. (B) Western blot of media from *Fusobacterium* growths shows that while truncated, FplA is not released into the media and remains associated with the bacteria. (C) Analysis of FplA expression during early exponential phase growth (OD_600_ = 0.2) reveals that strains 10953, which is cleaved in mid-exponential and stationary phase, is still in full-length state with a portion beginning to be cleaved. (D) Sequence alignment reveals that all FplA sequences from cleaved strains contain a highly charged motif at the PLA 1/POTRA hinge region as a potential site for an unidentified protease, with the exception being the non-cleaved FplA proteins from *F. nucleatum* strains 23726 and 25586, which contain a drastically different neutral motif.

While the FplA amino acid sequences from the seven tested strains are highly similar (>95% identity), we identified two regions in *F. nucleatum* 23726 and *F. nucleatum* 25586 at the intersection of the end of the N-terminal extension and just before the end of the PLA_1_ domain, which could correspond to potential protease processing sites (**Fig. 6D**, **Fig. S6**). The suspected cleavage site in *F. nucleatum* 23726 and *F. nucleatum* 25586 flanking the PLA_1_ domain is switched from a highly-charged motif (consensus sequence: KNIEDKKEKF), to a more neutral motif (consensus sequence: KFVTNSDAKI) that could be more protease resistant, resulting in retention of the full-length protein. In addition, to arrive at the 25 kDa product seen in five strains, a second cleavage event could occur at the end of the N-terminal extension, as strains 23726 and 25586 differ in this region by substitution of an alanine for charged and polar residues (**Fig. S6)**.

### FplA binds phosphoinositide signaling lipids with high affinity and could play a role in host colonization and altered signaling

We first demonstrated that FplA is a potent phospholipase with PLA_1_ activity (**Fig. 7A**) using artificial fluorogenic substrates. Next, we tested FplA for binding to lipids found in human cells and found that it preferentially binds to human phosphoinositides, as was previously seen when characterizing the homologous enzyme PlpD from *P. aeruginosa*^29^ (**Fig. S7**). Upon incubation with a more diverse and freshly-prepared library of PIs, FplA was found to preferentially bind to PI(4,5)p_2_, and with even stronger affinity to PI(3,5)p_2_, and PI(3,4,5)p_3_ lipids (**Fig. 7b**). This is consistent with structurally homologous enzymes binding PIs, and implicates a role for this enzyme in an intracellular environment.

**Fig 7.**
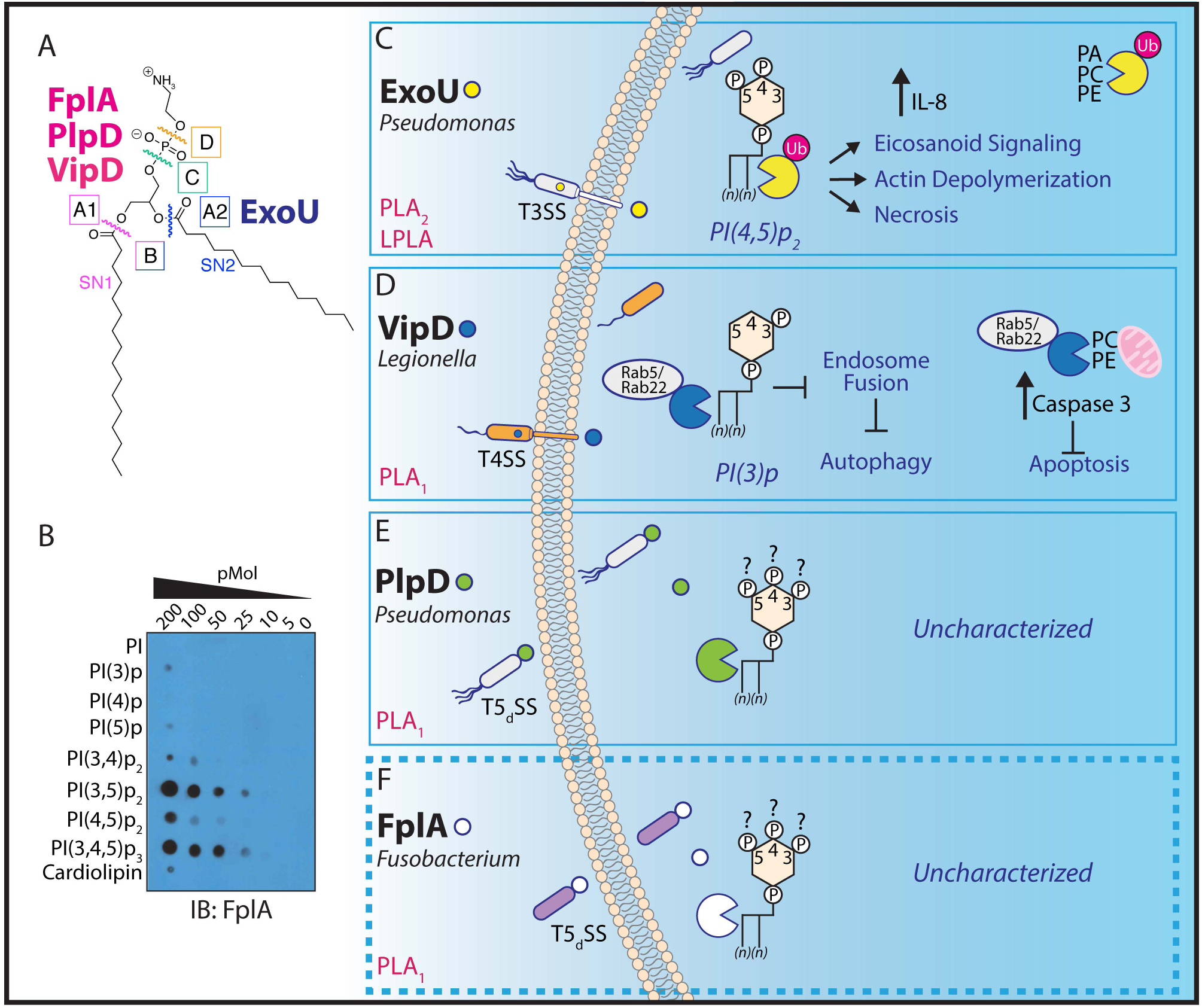
Overview of select bacteria phospholipase A enzymes and the role they play in intracellular processes. (A) Overview of phospholipase enzyme classes and the bonds they cleave within a phospholipid. (B) FplA_20-431_ binds with high affinity to phosphoinositide lipids that are critical for multiple cellular processes in a human host. (C-F) Bacterial phospholipases are confirmed virulence factors that play a major role in colonization of the host and evasion of autophagy and subsequent clearance. While the role of ExoU (T3SS) and VipD (T4SS) have been well characterized, the role of the TSdSS PLA 1 enzymes PlpD from *P. aeruginosa* and FplA from *F. nucleatum* remain to be determined.

## DISCUSSION

*F. nucleatum* is an emerging pathogen with an increasingly identified role in the onset and progression of colorectal cancer^8,10–13^. Because of this as well as other strong connections with pathogenesis, tools are needed to probe the molecular mechanisms used by this bacterium during host colonization and subsequent infection. Seminal work by other groups has shown a repertoire of both small (FadA, ∼15 kDa) and large (Fap2, >300 kDa, Type Va secreted) adhesins that are critical for interaction with the host to initiate entry into cells; in turn critical for the onset of inflammation^2,17^. Upon entry into host cells, very little has been reported about how *F. nucleatum* is able to establish an intracellular niche. We set out to probe the role of a potential virulence factor that we predicted to have phospholipase activity, thereby providing a potential mechanism for colonization and subversion of the host mechanisms of bacterial clearance. We characterized the gene FN1704, which we have renamed *fplA* for *Fusobacterium* phospholipase autotransporter (FplA). Our *in vitro* studies were focused on identifying tools and methods to characterize Type Vd secreted autotransporters to determine their role in virulence in a diverse set of Gram-negative bacteria; many such autotransporters have been identified in intracellular pathogens^28^. We created a *F. nucleatum* 23726 Δ*fplA* strain which will allow us to next probe the role of this enzyme through the first *in vivo* studies of Type Vd autotransporter phospholipases in infection. Our analyses indicate that deletion of the *fplA* gene from *F. nucleatum* does not alter growth or cell size and shape under laboratory growth conditions, adding to our hypothesis that FplA is a virulence factor and not a bacterial maintenance protein. Considering both PlpD and FplA are Type Vd autotransporters that bind human phosphoinositides, they have the potential to share the same *in vivo* role as the structurally similar phospholipases ExoU and VipD, which have been characterized as intracellular virulence factors involved in cleavage of PI(4,5)p2 and PI(3)p respectively. Hydrolysis of these lipids result in modulation of the host by inducing actin depolymerization, cell death, and subversion of host responses to bacterial infection by blocking autophagy and apoptosis^38,40–42^.

Determining that different *F. nucleatum* strains express mature FplA proteins of varying molecular weights was a surprising result that made us question which form of the enzyme may be involved in virulence. Because of the genetic intractability of *Fusobacterium* spp., we have not been able to delete copies of *fplA* in strains that we predicted to have a truncated yet surface-associated version of the protein. The development of more robust genetic systems for *Fusobacterium* has the potential to open doors to fill a critical knowledge gap in the role of Type Vd secretion in a variety of clinically isolated *F. nucleatum* strains.

While our focus has been on the role of FplA induced changes in intracellular signaling, it still remains to be determined if this enzyme plays a role in other processes by cleaving additional host lipids. Support for this comes from results showing ExoU also cleaves PA, PC, and PE^38,54^, and VipD is able to cleave PC and PE^32^. Our initial results showed that FplA does not bind with high affinity to PA, PC, and PE, but these results do not rule out potential cleavage of these lipids in experiments that better simulate an environment found in infection. FplA could be involved in cleaving lipids in a mucous-rich environment, as this bacterium is found in the gut and virulent strains have been isolated from patients with inflammatory bowel disease^15^. To add to the role of bacterial phospholipases cleaving lipids found in structural membranes, ExoU plays a major role in *P. aeruginosa* entry into the bloodstream upon leaving the lungs^55^, and strains lacking ExoU are far less virulent and are cleared more efficiently in mouse models of pneumonia^56,57^. Since cases of *Fusobacterium* bacteremia are frequently documented (*F. nucleatum* comprises 61% of cases)^58^, and a wide array of bodily locations have been reported for *F. nucleatum* infections (brain^59^, liver^60^, lungs^5^, heart^61^), it will be critical to use our characterized chemical and biochemical tools, including an *fplA* deletion strain, to test the role of this enzyme in the previously established hematogenous spread^2^.

Outside of the Type III, IV, and V secreted phospholipases A1 and A2, *Vibrio cholerae* also produces a large Type I secreted multifunctional-autoprocessing repeats-in-toxin (MARTX) protein characterized as a PLA1 enzyme that selectively cleaves PI(3)p and upon expression in mammalian cells, reduces intracellular PI(3)p levels and inhibits endosomal and autophagic pathways^62^. In addition, MARTX from *Vibrio vulnificus* is necessary for epithelial barrier disruption and intestinal spread^63^. Unique to MARTX when compared to FplA, PlpD, VipD, and ExoU, is the use of a catalytic triad (Ser, Asp, His) instead of a dyad (Ser, Asp)^62^. To add to the importance of bacterial enzymes that alter phosphoinositides, the *Mycobacterium tuberculosis* enzyme SapM is a PI(3)p-specific phosphatase that depletes phagosomes of this signaling lipid, resulting in the inability of the infected cell to form mature phagolysosomes^64^. *Legionella pneumophila* was recently found to use the phosphoinositide kinase LepB to convert PI(3)p to PI(3,4)p_2_, and subsequently SidF, a phosphoinositide phosphatase converts PI(3,4)p_2_ to PI(4)p, which is an important docking molecule for multiple *L. pneumophila* effectors to label the *Legionella*-containing vacuole inside host cells^65^.

As there are an impressive number of phosphoinositide modulating enzymes secreted by bacteria to alter host signaling and induce colonization, it will be important to develop a robust set of chemical and molecular tools to determine the role of Type Vd surface bound or secreted PLA_1_ enzymes in virulence and intracellular survival. FplA is the lone Type Vd PLA_1_ enzyme found in *F. nucleatum*, and we believe, based on our biochemical analysis and multiple previously characterized functions of phospholipases in pathogenic bacteria, FplA has the potential to be critical for the intracellular survival and pro-oncogenic signaling by this emerging pathogen.

## MATERIALS AND METHODS

### Bacterial strains, growth conditions, and plasmids

Unless otherwise indicated, *E. coli* strains were grown in LB at 37°C aerobically, and *F. nucleatum* strains were grown in CBHK (Columbia Broth, hemin (5 μg/mL) and menadione (0.5 μg/mL)) at 37°C in an anaerobic chamber (90% N_2_, 5% CO_2_, 5% H_2_). For taxonomy verification of *Fusobacterium*, PCR amplification of a 1502bp region of the 16S rRNA gene sequence was carried out using the universal primers U8F and U1510R (**Table S3**) as previously described^66^. The primers were used at a concentration of 20 μM with 1-2 μL of extracted genomic DNA as the template. The reaction conditions were: 95°C 3 min, (98°C 20 s, 50°C 15 s, 72°C 1 min) x 35 and 72°C 5 min. The quality of the amplicons was determined by agarose gel electrophoresis, and amplicons were then purified using the EZ-10 Spin Column PCR Products Purification Kit (BioBasic, Canada), quantified on the NanoDrop® ND-8000 (Thermofisher; Burlington, ON) and sent for Sanger sequence analysis following a BigDye® Terminator v.3.1 cycle sequencing PCR (Thermofisher; Burlington, ON) amplification. Sanger sequencing was carried out at the Advanced Analysis Center at the University of Guelph. Obtained DNA sequences were compared to the GenBank database (NCBI) using BLASTn. Where appropriate, antibiotics were added at the indicated concentrations: carbenicillin 100 μg/mL; thiamphenicol 5 μg/mL (CBHK plates), 2.5 μg/mL (CBHK liquid).

### Bioinformatic analysis of *fplA* in multiple *Fusobacterium* strains

The genome sequence of *F. nucleatum* strain ATCC 25586 (GenBank accession NC_003454.1) was used to predict all open reading frames using the Prodigal Bacterial Gene Prediction Server^67^. An open reading frame encoding for a 760 amino acid protein was identified using a HMMER model built from a seed alignment of the PFAM (EMBL-EBI website) patatin family (PF01734) and the stand alone HMMER 3.1 software package^68^. The identified gene contained an N-terminal patatin domain conferring phospholipase activity and a C-terminal bacterial surface antigen domain (PFAM: PF01103) that encodes for an outer membrane beta barrel domain. Cross referencing revealed this gene is FN1704 in *F. nucleatum* ATCC 25586, which was incorrectly predicted to be a serine protease in both the KEGG and Uniprot databases. The same method was used to search multiple *Fusobacterium* genomes resulting in the identification of only one protein with this structure in each strain. A PSI-BLAST search using FplA returned a close match to the *Pseudomonas aeruginosa* protein PlpD, which was previously characterized as a class A1 phospholipase and labeled as the first in a new class of type Vd autotransporters^28,29^. Alignment of FplA proteins from seven strains of *Fusobacterium* shown in **Fig. S6** was performed using Geneious version 9.0.2^69^.

### Structure prediction to identify domain boundaries and catalytic residues in FplA

Structure prediction was performed using the FplA sequence from *F. nucleatum* strain 25586 and the SWISS-MODEL Workspace^70^. Results showed a close match of the N-terminal phospholipase domain to PlpD from *Pseudomonas aeruginosa* (PDB: 5FYA) and the C-terminal POTRA and beta barrel domains to BamA from *Haemophilus ducreyi* (PDB: 4K3C) (**Fig. 1B, Fig. S1B**). A composite predicted structure was assembled using the predicted phospholipase, POTRA, and beta barrel domain, which has the phospholipase domain exposed on the surface of the bacteria (**Fig. S1**), which we confirmed biochemically as a recombinant protein in *E. coli* and a native protein in *F. nucleatum*. In addition, the modeled FplA phospholipase domain was aligned with ExoU (PDB: 4AKX) and VipD (PDB: 4AKF) (**Fig. S2A-B**). Active site residues in FplA were identified as S98 and D243, and these were verified by multiple enzymatic and chemical biology methods presented in **Fig. 2** and **Fig. 3**. In close proximity to the active site is the oxyanion hole comprised of three consecutive glycine residues (G69, G70, G71). Graphical representations and alignments of all predicted structures were created using PyMOL Molecular Graphics System, Version 1.7.3 Schrödinger, LLC.

### Cloning of FplA Constructs for Expression in *E. coli*

All primers were ordered from IDT DNA and all plasmids and bacterial strains either used or created for these studies are described in **Table S1** (Bacterial Strains), **Table S2** (Plasmids), and **Table S3** (Primers). All restriction enzymes and T4 DNA Ligase, and Antarctic Phosphatase were from NEB (MA, USA). DNA purifications kits were from BioBasic (Markham, ON). Genomic DNA for *F. nucleatum* ATCC 25586 was purchased from ATCC (VA, USA) and used to create all recombinant FplA constructs for expression described herein. pET16b was used as the base expression vector for *E. coli* expression of FplA constructs. All constructs were created by using 50 ng of genomic DNA as a template, followed by PCR amplification with primers for each construct described in **Table S3** using Q5 High-Fidelity Polymerase (NEB, USA) and a ProFlex PCR System (Applied Biosystems, USA) under the following conditions: 98°C 2 min, (98°C 20 s, 50-62°C 20 s, 68°C 1-4 min) x 6 cycles for 50, 53, 56, 59, 62°C (30 cycles total), and 72°C 5 min. PCR products were then spin column purified and digested overnight at 37°C with restriction enzymes described in **Table S3**. Digested PCR products were spin column purified and ligated by T4 DNA ligase into pET16b vector that had been restriction enzyme and Antarctic Phosphatase treated according to the manufacturer′s recommended protocol in a 20 μl final volume for 1 hour at 26°C. 5 μl of ligations were transformed into Mix & Go! (Zymo Research, USA) competent *E. coli* and plated on LB 100 μg/ml carbenicillin (ampicillin), followed by verification of positive clones by restriction digest analysis using purified plasmid. Positive clones were then transformed into LOBSTR RIL^71^ *E. coli* cells for protein expression.

Specifically, pDJSVT84 (FplA_20-350_), pDJSVT43 (FplA_20-431_), pDJSVT85 (FplA_60-350_), and pDJSVT82 (FplA_60-431_) all produce proteins with a C-terminal 6x-Histidine tag and are expressed in the cytoplasm because these constructs lack the N-terminal signal sequence used to export FplA through the Sec apparatus in *F. nucleatum*. pDJSVT60 (FplA_20-431_ S98A) and pDJSVT61 (FplA_20-431_ D243A) were created by using pDJSVT43 as a template for Quikchange mutagenesis PCR. Verification of mutants and all clones was performed by Sanger sequencing (Genewiz, USA). To facilitate the export of FplA to the surface of *E. coli*, a new inducible expression vector was created using pET16b as the backbone by incorporating the signal sequence from the *E. coli* protein OmpA (residues 1-27). In addition, this expression vector (pDJSVT86) contains an N-terminal 6x-Histidine tag that remains on the expressed protein after residues 1-21 from OmpA are cleaved in the periplasm. This effectively creates an inducible vector for the expression of periplasmic and outer membrane proteins in *E. coli* that was customized with GC rich restriction sites (*NotI, KpnI, XhoI*) to facilitate enhanced cloning of AT rich (74%) genomes such as *F. nucleatum*. Using the pDJSVT86 expression vector, pDJSVT88 (OmpA_1-27_, 6xHis, FplA_20-760_) was created and shows efficient export of enzymatically active, full-length FplA to the surface of *E. coli* (**Fig. 4**).

### FplA Protein Expression and Purification

Briefly, all FplA construct in LOBSTR RIL^71^ *E. coli* cells were grown in Studier auto-induction media^72^ (ZYP-5052, 0.05% glucose, 0.5% lactose, 0.5% glycerol) at 37 °C, 250 rpm shaking, and harvested at 20 hours post inoculation by pelleting at 5 kG for 15 minutes at 4°C. Pellets were weighed and resuspended in lysis buffer (20 mM tris pH 7.5, 20 mM imidazole, 400 mM NaCl, 0.1 % BOG, 1 mM PMSF) at 10 mL/gram of cell pellet. Bacteria were lysed by using 5 passes on an EmulsiFlex-C3 (Avestin, Germany), followed by removal of insoluble material and unlysed cells by pelleting at 15 kG for 15 minutes at 4°C. The resulting supernatant containing 6xHis-tagged FplA constructs were gently stirred with 5 mL of NiCl_2_ charged chelating sepharose beads (GE Healthcare, USA) for 30 minutes at 4°C, followed by washing with 200 mL of wash buffer (20 mM Tris pH 7.5, 50 mM Imidazole, 400 mM NaCl, 0.1% BOG). After washing, FplA was eluted in 10 mL of elution buffer: (20 mM Tris pH 7.5, 250 mM Imidazole, 50 mM NaCl, 0.1% BOG). This protein was directly applied to a HiTrap Q FP anion exchange column (FplA construct theoretical PIs: 5.91-6.34) and purified on an ÄKTA pure system (GE Healthcare, USA) using a linear gradient between Buffer A (20 mM Tris, pH 8, 50 mM NaCl, 0.025% BOG) and Buffer B (20 mM Tris, pH 8, 1 M NaCl, 0.025% BOG). Fractions containing FplA as determined by SDS-PAGE analysis were pooled and further purified on a High-prep 16/60 Sephacryl S-200 HR size exclusion column (GE Healthcare, USA) in 20 mM Tris pH 7.5, 150 mM NaCl, 10% glycerol. Protein concentrations were determined using a Qubit fluorimeter and BCA assays according to the manufacturer′s recommended protocol. Protein purity was determined using ClearPage 4-20% gradient gels (CBS Scientific, USA) and determined to be greater than 95% pure for all constructs.

### Antibody Production and Western Blotting to Detect FplA

Purified FplA_20-431_ was used to create a polyclonal antibody in rabbits (New England Peptide, USA). To purify the antibody, FplA_20-431_ was coupled to CNBr-Activated Sepharose (Bioworld, USA) and Anti-FplA_20-431_ antisera adjusted to pH 8.0 with 20 mM Tris-HCl was passed through the column to bind FplA_20-431_ antibodies, followed by extensive washing in phosphate buffered saline (PBS) and elution in 2.7 mL of 100 mM Glycine, pH 2.8. To the eluted antibodies, 0.3 mL of 1M Tris-HCl pH 8.5 was added, for a final storage buffer of (10 mM Glycine, 100 mM Tris-HCl, pH 8.5).

For western blot detection of FplA, proteins were separated by SDS PAGE gels run at 210V for 60 minutes, followed by transferring proteins to PVDF membranes in transfer buffer (25 mM Tris, 190 mM Glycine, 20% methanol, pH 8.3) at 80V for 60 minutes. Post-transfer, membranes were blocked in 20 mL of TBST (20 mM Tris, 150 mM NaCl, 0.1% Tween 20) with 3% BSA for 15 hours at 4°C. After blocking, the membranes were incubated with rabbit anti-FplA antibody (1:10,000 for pure proteins, 1:2,500-1:1000 whole cells or lysates) in TBST 3% BSA for 1 hour (70 rpm shaking, 26°C). After incubating with the primary antibody the membrane was washed with TBST, followed by incubation with goat anti-rabbit-HRP secondary antibody (Cell Signaling, USA) at 1:10,000 dilution in TBST 3% BSA for 30 minutes (70 rpm shaking, 26°C). After the secondary antibody incubation, the membrane was washed in TBST, followed by incubation with ECL-Plus blotting reagents (Pierce, USA) and visualization using Lucent Blue X-ray film (Advansta, USA) developed on an SRX-101A medical film processor (Konica, Japan).

### Development of an *F. nucleatum* 23726 Δ*fplA* strain

Single-crossover homologous recombination gene knockouts *F. nucleatum* 23726 have been previously reported, although like with all *Fusobacterium* mutagenesis strategies, efficiencies are quite low. Based on a previous method^52^, we created an integration plasmid that will not replicate in *F. nucleatum*, therefore only producing antibiotic resistant colonies for strains that incorporate the plasmid directly into the chromosome in the gene of interest during transformation and outgrowth. A central 1000bp region in the FN1704 (*fplA)* gene in *F. nucleatum* 23726 was amplified from genomic DNA by PCR, digested with *EcoRI* and *SpeI*, and ligated into pJIR750 that was digested with the same enzymes and subsequently treated with Antarctic phosphatase. The ligation was transformed into Mix & Go! competent *E. coli*, and plated on LB 10 μg/mL chloramphenicol, followed by selection of colonies, purification of plasmid DNA, and verification of positive clones by restriction digest analysis. A single positive clone was selected for all future studies, and DNA was initially purified by spin column (BioBasic, Canada), followed by additional purification of the DNA using glycogen and methanol precipitation, followed by resuspension in sterile deionized H_2_O.

*F. nucleatum* 23726 was made competent by growing a 5 mL culture to mid-log phase (OD_600_ = 0.4) followed by spinning down cells at 14k G for 3 minutes, removal of media, and five successive 1 mL washes with ice cold 10% glycerol in diH_2_O. Cells were then resuspened in a final volume of 100 μL of ice cold 10% glycerol (Final OD_600_= ∼20). Bacteria were transferred to cold 1 mm electroporation cuvettes (Genesee, USA) and 0.5-2.0 μg ([ ] > 500 ng/μL) of pDJSVT100 plasmid was added immediately before electroporating at 2.0 kV (20 kV/cm), 50 μF, 129 OHMs, using an Electro Cell Manipulator 600 (BTX, USA). To the cuvette, 1 mL of recovery media (CBHK, 1 mM MgCl_2_) was added, and immediately transferred by syringe into a sterile, anaerobic tube via septum for incubation at 37°C for 20 hours with no shaking. Post outgrowth, cells were spun down at 14 kG for 3 minutes, media removed, and resuspended in 0.1 mL recovery media, followed by plating on CBHK plates with 5 μg/mL thiamphenicol and incubation in an anaerobic 37°C incubator for two days for colony growth. ∼ 5 colonies/μg of DNA were achieved, and the *fplA* gene knockout was verified by PCR specific to the chromosome and *catP* gene that was incorporated into the genome by the pDJSVT100 KO plasmid (Primers, **Table S3)**. In addition, western blots were used to confirm a loss of FplA protein expression (**Fig. 5-6**).

### Enzymatic assay design, data collection, and FplA kinetics

Initial tests for FplA enzymatic activity were run using the EnzChek Phospholipase A1 and EnzChek Phospholipase A2 assay kits (ThermoFisher, USA) at 1 μM and 10 μM FplA_20-431_ using the manufacturer′s protocol (**Fig. S3A-B**). These assays showed that FplA has PLA_1_, but not PLA_2_ activity, which is consistent with data reported for the homologous enzyme PlpD. We then went on to further characterize its activity by developing a continuous kinetic assay using the PLA_1_ specific substrate PED-A1 (ThermoFisher, USA) and determined the full kinetic parameters of FplA with this substrate as reported in **Fig. 2** and **Fig. S3**. In detail, FplA was used at 1 nM in the reaction and substrate (10 mM stock in 100% DMSO) dilutions (0-10 μM) and reactions were carried out in reaction buffer (50 mM Tris pH 8.5, 50 mM NaCl, 0.025% BOG). All samples including controls contained equal concentrations of DMSO. Reactions were run at 26°C for 30 minutes with 3 seconds of shaking in between continuous fluorescent monitoring (Ex = 488 nm, Em = 530 nm) every 2 minutes on a SpectraMax M5^e^ plate reader (Molecular Devices, USA). Relative fluorescence units measured upon cleavage of substrate ester bonds and release of the acyl chain were converted to the concentration of product (BODIPY® FL C5) created by establishing a standard curve using pure BODIPY® FL C5 (ThermoFisher, USA). In all enzymatic reactions, controls containing no protein were run and the values were subtracted from the reactions containing protein during analysis.

We then developed a continuous fluorescent assay to characterize the phospholipase activity of FplA using the general lipase substrates 4-Methylumbelliferyl Butyrate (4-MuB) and 4-Methylumbelliferyl heptanoate (4-MuH) (Santa Cruz Biotechnology, USA). In detail, FplA was used at 1 nM in the reaction and substrate (50 mM stock in 100% DMSO) dilutions (0-200 μM) and reactions were carried out in reaction buffer (50mM Tris pH 8.5, 50mM NaCl, 0.025% BOG). All samples including controls contained equal concentrations of DMSO (0.4%). Reactions were run at 26°C for 30 minutes with 3 seconds of shaking in between continuous fluorescent monitoring (Ex = 360 nm, Em 449 nm) every 2 minutes on a SpectraMax M5^e^ plate reader. Relative fluorescence units measured upon cleavage of substrate ester bonds and release of the acyl chain were converted to the concentration of product (4-Methylumbeliferone, 4-Mu) created by establishing a standard curve using pure 4-Mu (Sigma Aldrich, USA).

The steady-state kinetic parameters for each substrate were determined using GraphPad Prism version 6 (Graphpad Software, USA) by fitting the initial rate data (n=2) to the Michaelis-Menten equation (**Equation 1**):

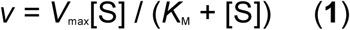

to obtain the values reported in **Fig. 2** and **Fig. S3**.

### Characterization of FplA inhibitors

We set out to characterize inhibitors that we could use as effective tools to test the role of FplA both *in vitro* and potentially *in vivo* by IC_50_ assays using a variety of inhibitor classes. Inhibitors shown in **Fig. 3** and **Fig. S4** were chosen based on their previous classification as inhibitors of a diverse set of phospholipase enzymes: Methylarachidonyl fluorophosphonate (MAFP), PLA^73^_2_; Arachidonyl Trifluoromethyl Ketone (ATFMK), cPLA_2_, iPLA^74^_2_; Isopropyl Dodec-11-Enylfluorophosphonate (IDEFP), fatty acid amide hydrolase^75^; Palmityl Trifluoromethyl Ketone (PTFMK), cPLA_2_, iPLA_2_^76^; ML-211, LYPLA1, LYPLA2^77^; Isopropyl Dodecylfluorophosphonate (IDFP), fatty acid amide hyrolase, monoacylglycerol lipase^78^; LY311727, sPLA_2_^45^; Manoalide, sPLA2, PLC^48,79^. All inhibitors were purchased from Cayman Chemical, USA.

For potent inhibitors, 0-25 μM concentrations were used in assays, and for compounds found to not inhibit efficiently, the concentration range was 0-100 μM. Inhibitors were diluted into reaction buffer (50 mM Tris pH 8.5, 50 mM NaCl, 0.025% BOG) containing 10 μM 4-MuH. To initiate the reaction, 1 nM final FplA_20-431_ was added and reactions were run at 26°C for 30 minutes with 3 seconds of shaking in between continuous fluorescent monitoring (Ex = 360 nm, Em 449 nm) every 2 minutes on a SpectraMax M5^e^ plate reader. Raw data (n=2) for each reaction were analyzed in GraphPad Prism using a log(inhibitor) vs. response using variable slope and a least squares (ordinary) fit model.

### Use of fluorescent chemical probes to label and detect FplA

Purified recombinant FplA constructs or WT FplA from *F. nucleatum* strains were visualized using an ActivX TAMRA-FP probe (ThermoFisher, USA). This probe only binds to proteins with activated serine residues. For purified recombinant proteins, 5 μg of purified protein was incubated with either 100 μM of methylarachidonyl fluorophosphonate (MAFP) or PBS for 1 hour. Following pre-incubation with MAFP or PBS, 1 μM ActivX TAMRA-FP probe was added to the protein and incubated for 20 minutes at 26°C followed by the addition SDS-PAGE running buffer to stop the reaction. 500 ng of protein was run on an SDS-PAGE gel at 210V for 60 minutes, followed by transferring proteins to PVDF membranes in transfer buffer (25 mM Tris, 190 mM Glycine, 20% methanol, pH 8.3) at 80V for 60 minutes. Fluorescent proteins were visualized using a G:Box XX6 system (SynGene, USA) using the TAMRA fluorescence filter.

For the detection of FplA in *F. nucleatum* whole cell mixtures, 5 ml of *F. nucleatum* 23726 or *F. nucleatum* 23726 Δ*fplA* cells at OD_600_ = 0.2 were pelleted, washed, and resuspended in 100 μL of PBS. ActivX TAMRA-FP was added at a final concentration of 2 μM and incubated at 26°C for 20 minutes, followed by the addition of SDS buffer. 10 uL of this reaction (lysate from ∼4.2 x 10^8^ bacteria) was run per well on an SDS-PAGE gel at 210V for 60 minutes. Gels were then imaged on a Typhoon Trio (GE Healthcare, USA) using the TAMRA filter setting.

### Detection of FplA on the surface of *E. coli* by microscopy, enzymatic activity, and Proteinase K treatment

Using the expression vector pDJSVT86 that is described in the cloning and expression section above, we cloned FplA_20-760_ into the vector at the 3′ end of the OmpA_1-27_-6xHis signal sequence (pDJSVT88). This construct was expressed in LOBSTR RIL^71^ *E. coli* in Studier autoinduction media at 37°C for 20 hours with 250 RPM shaking. The empty vector pDJSVT86 was used as a negative control for FplA expression for both microscopy and enzymatic assays.

For microscopy, stationary phase bacteria from overnight expressions were washed in PBS pH 7.5, 0.2% gelatin and spun down at 5 kG for 5 minutes, followed by resuspending the bacteria at an OD_600_=0.2. To the bacteria, a final 3.2% paraformaldehyde was added for 15 minutes at 26°C for fixation, followed by washing in PBS pH 7.5, 0.2% gelatin. 500 μL of fixed bacteria were then added on top of a polylysine coated coverslip in a 6 well plates, and 2 mL of PBS was added for a final volume of 2.5 mL. Bacteria were then spun down onto the coverslips at 2,000x G for 10 minutes. Washed coverslips were submerged in 300 μL of PBS pH 7.5, 0.2% gelatin containing a 1:100 dilution of the anti-FplA antibody and incubated for 20 hours at 26°C with light shaking. Coverslips were washed again in PBS pH 7.5, 0.2% gelatin and then incubated in the same buffer containing an anti-rabbit Alexa Fluor 488 conjugated secondary antibody for 30 minutes at 26°C. Washed coverslips were mounted with Cytoseal 60 (ThermoFisher, USA) and visualized by brightfield and fluorescence microscopy using the GFP channel on an EVOS FL microscope (Life Technologies, USA).

For enzymatic the enzymatic activity assay, stationary phase bacteria from overnight expressions were washed in PBS pH 7.5 and spun down at 5 kG for 5 minutes, followed by resuspending the bacteria at an OD_600_=0.2 in PBS pH 7.5. Bacterial samples were incubated with 10 μM MAFP or PBS pH 7.5 at RT for 60 minutes at 26°C, followed by washing in PBS pH 7.5 and resuspension to the original OD_600_=0.2 (2 x 10^8^ CFU/mL in enzymatic assay buffer (50 mM Tris pH 8.5, 50 mM NaCl, 0.025% BOG). 2 x 10^6^ bacteria were then added to reaction wells containing 10 μM 4-MuH fluorescent lipase substrate (Ex = 360 nm, Em 449 nm), followed by incubation at 37°C for 30 minutes and detection of lipid cleavage and product formation with a Spectramax M5^e^ as seen in **Fig. 4B**. Activity was plotted as fluorescence units and Statistical analysis was performed using a multiple comparison analysis by one-way ANOVA in GraphPad Prism.

To further validate the translocation of the PLA_1_ domain of FplA to the surface of *E. c*oli, the non-specific and membrane impenetrable enzyme proteinase K (PK) was used to cleave FplA in a dose dependent manner. FplA expression was induced with 500 μM IPTG for four hours shaking at 37°C. Bacteria were washed in PBS and adjusted to an OD_600_ = 0.2 in PBS with 1 mM CaCl_2_ to activate PK. 100 μL of cells were added to tubes followed by the addition of 0, 100, 250, or 1000 nM PK and incubation at 26°C for 15 minutes. Reactions were then quenched with protease inhibitors (Roche, USA) and samples were separated by SDS-PAGE and transferred to PVDF for western blot analysis with an anti-FplA antibody. As a control, *E. coli* with the empty vector pDJSVT86 were analyzed for FplA expression and cleavage. In addition, GAPDH was used a load control, and also as a control to show PK was not digesting intracellular proteins.

### Lipid binding assays

Binding of FplA to various lipids was performed with commercially available lipids spotted on membranes, or by our laboratory spotting fresh lipids on blots.

For the first analysis, membrane lipid strips were purchased from Eschelon, Inc. The strips were blocked in 10 mL of TBST 3% BSA for 2 hours at 26°C with 70 rpm shaking. After blocking, lipid strips were incubated with TBST 3% BSA containing 50 μg/mL of the indicated FplA construct at 4°C for 15 hours. After incubation with FplA, lipid strips were washed with TBST and incubated with a 1:1000 dilution of rabbit anti-FplA antibody in 10 mL of TBST 3% BSA for 60 minutes at 26°C with 70 rpm shaking. Lipid strips were washed with TBST and incubated with a 1:2000 dilution of goat anti-rabbit IgG-HRP linked antibody (Cell Signaling, USA) in 10 mL of TBST 3% BSA for 30 minutes at 26°C with 70 rpm shaking. After secondary antibody incubation, the lipid strips were thoroughly washed in TBST, and ECL-Plus blotting reagents were added for visualization. The membranes were visualized using a G:Box XX6 system (SynGene, USA) (**Fig. S7**)

For a more detailed analysis of FplA binding to phosphoinositides, we purchased various phosphoinositides from Avanti Polar Lipids, and then spotted them onto PVDF at concentrations from 0-200 picomols (pMol). We tested FplA binding to PI, PI(3)p, PI(4)p, PI(5)p, PI(3,4)p_2_, PI(3,5)p_2_, PI(4,5)p_2_, PI(3,4,5)p_3_, and cardiolipin. All steps for analysis were the same as described above, except the membranes were visualized using Lucent Blue X-ray film developed on a SRX-101A medical film processor (**Fig. 7B**).

## ACKNOWLEDGEMENTS

We thank the following individual for help and guidance with these studies: S. Melville (Virginia Tech) for critical insight regarding bacterial mutagenesis and for providing the pJIR750 plasmid; C. Caswell (Virginia Tech) for critical conversations regarding bacterial genetics; W. Lewis (WUSTL) for help with the *Fusobacterium* electroporation protocol; M. Klemba (Virginia Tech) for reagents and critical phospholipase insights. Partial funding for this work was provided by Virginia Tech new faculty start-up funds to DJS. Partial funding for this work was provided through an Innovation grant to EA-V from the Canadian Cancer Society Research Institute.

---

**Fig S1.**
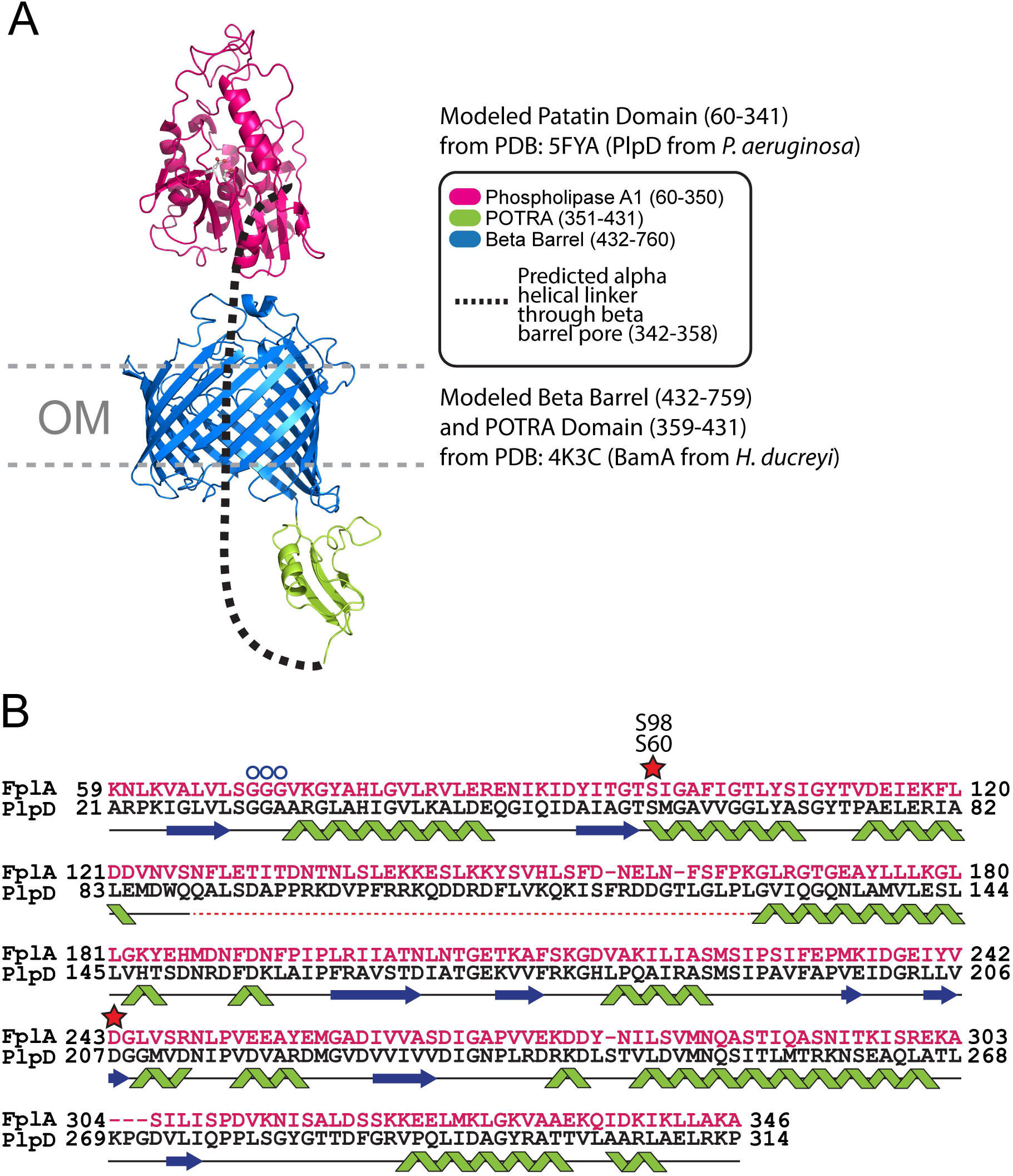
(A) Composite model of the predicted FplA structure in a bacterial outer membrane (OM) (B) Alignment of amino acids from PlpD and FplA from the predicted structures of the PLA_1_ domains.

**Fig S2.**
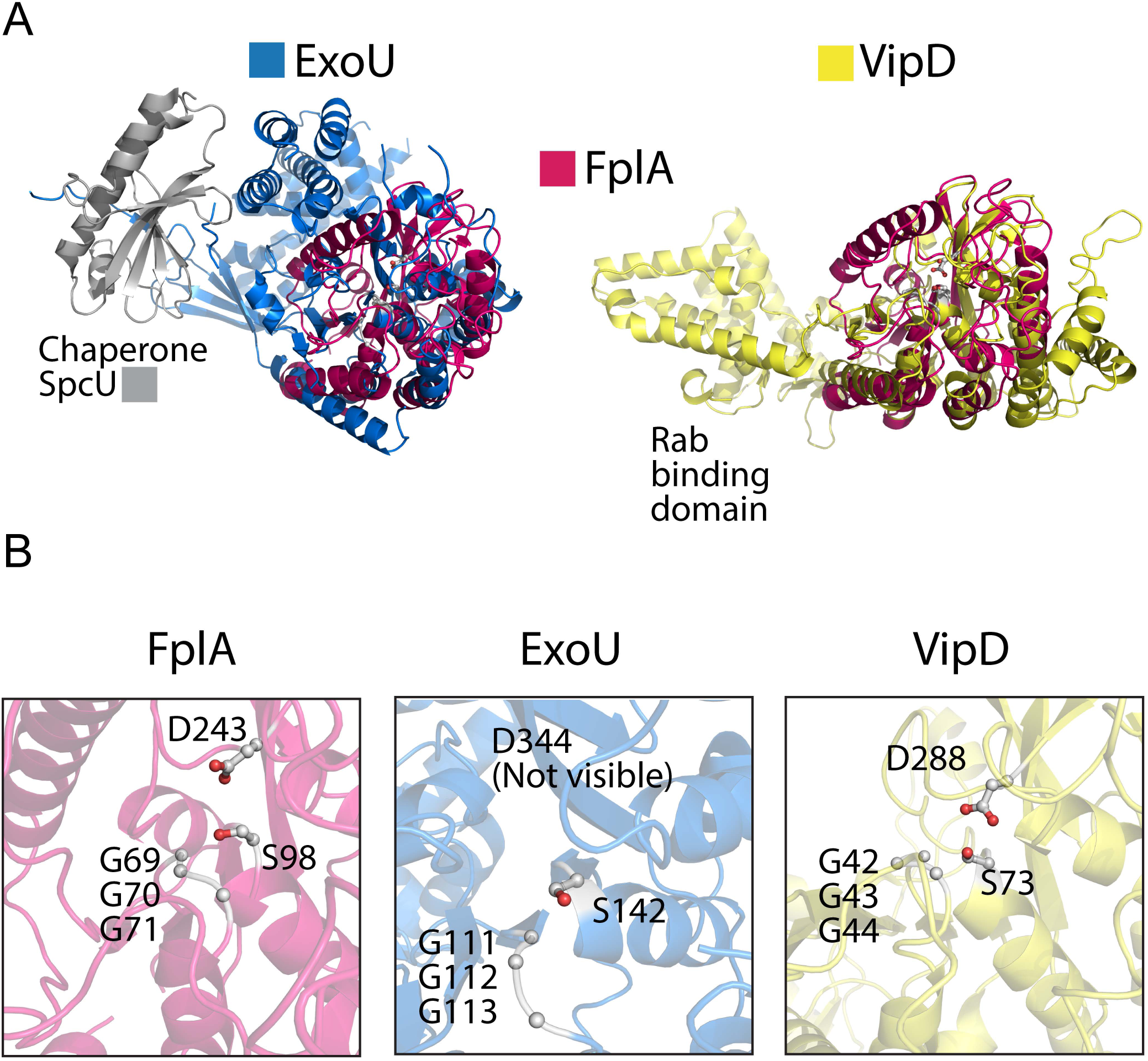
(A) Alignments of a predicted FplA structure (residues 60-431) with ExoU (PDB: 4AKX) and VipD (PDB: 4AKF). (B) Zoomed in view of active sites after alignment showing similar architectures and residue placement of the catalytic dyad (Ser, Asp) and oxyanion hole (Gly, Gly, Gly).

**Fig S3.**
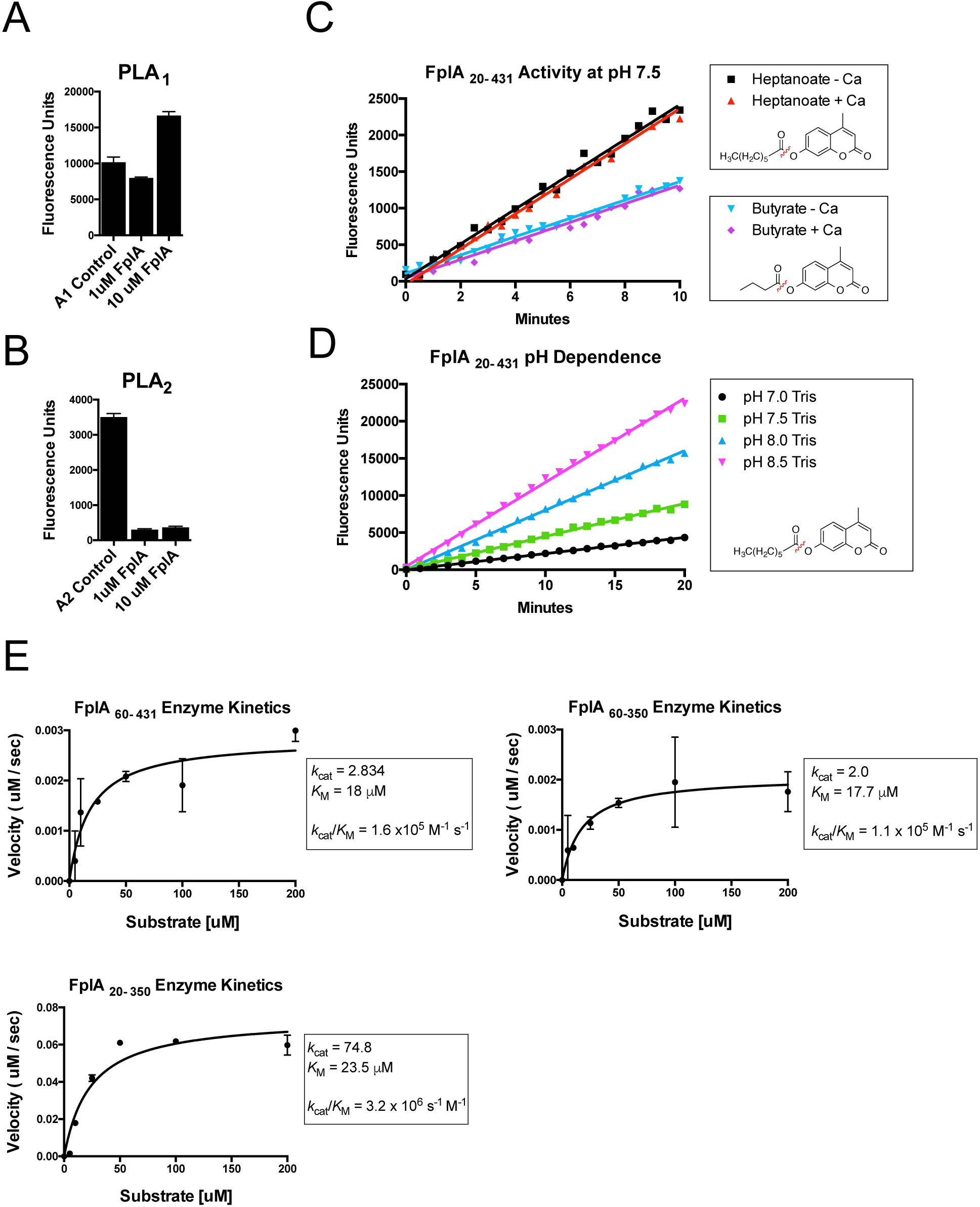
Eznymatic analysis of FplA_20-431•_ (A-B) Enzymatic assays show that FplA is a PLA_1_ specific enzyme with no PLA_2_ activity. (C) FplA_20-431_ does not need calcium for activity and is more active against the substrate 4-MuH than 4-MuB, indicating that longer acyl chains are critical for substrate binding. (D) pH dependent active of FplA_20-431_ using 4-MuH as a substate shows maximal activity at pH 8.5. (E) Eznyme kinetics and activity plots of FplA_60-431,_ FplA_60-350,_ and FplA_20-350_ with 4-MuH.

**Fig S4.**
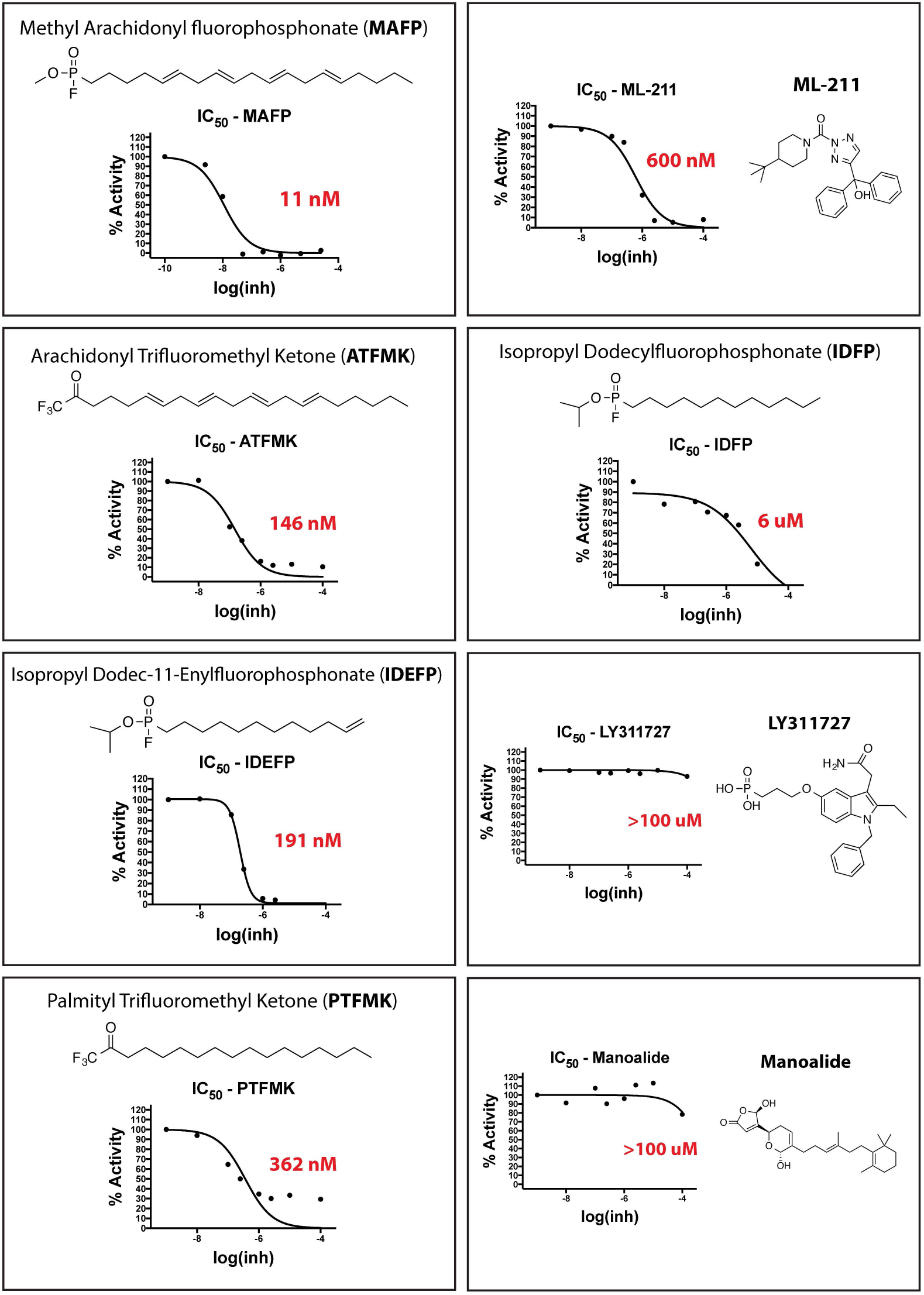
Analysis of multiple inhibitors previously shown to inhibit a diverse set of phospholipases. ICSO values equte to the concentration of inhibitor necessary to achieve 50% inhibition of FplA_20-431_ using 4-MuH as a substate.

**Fig S5.**
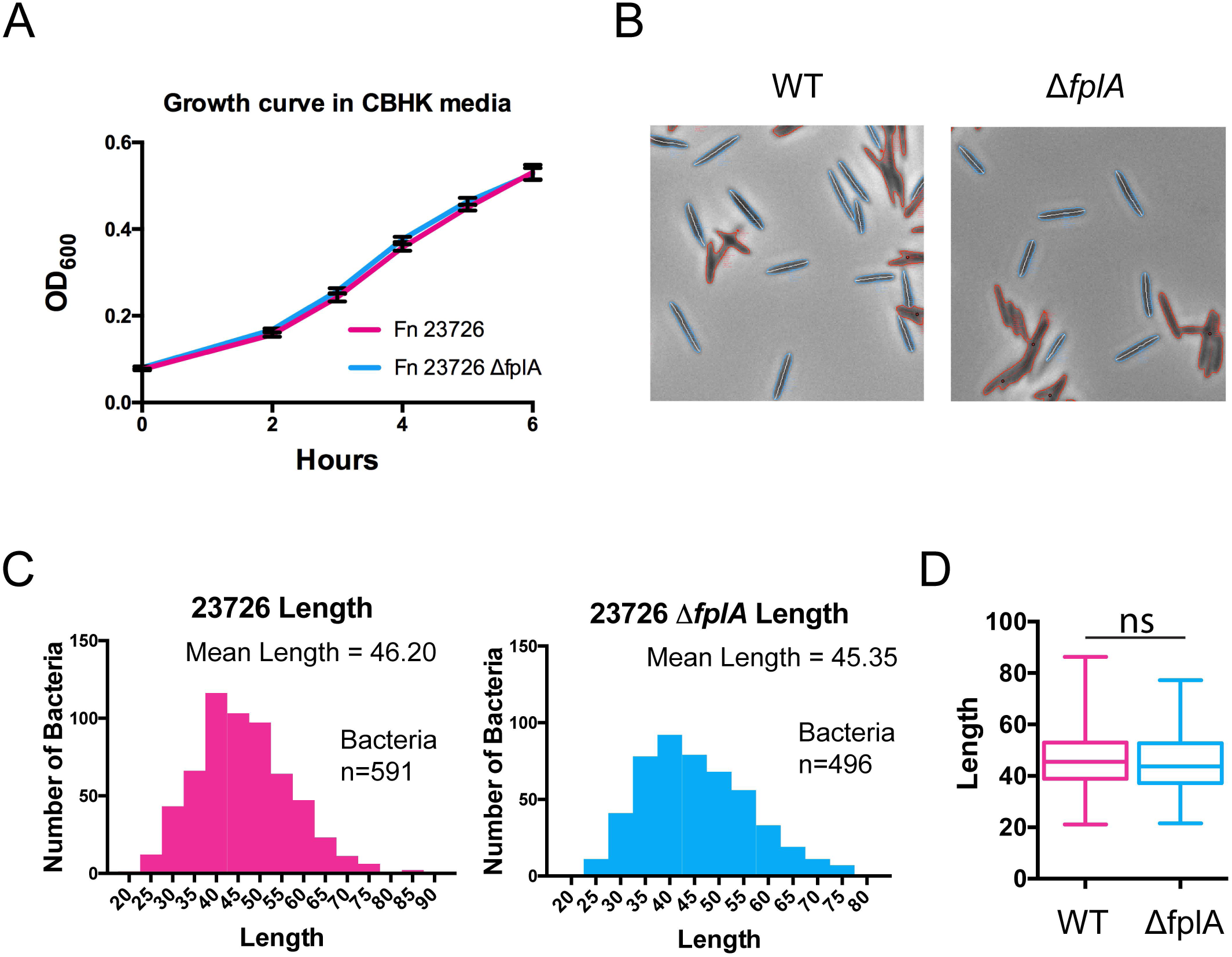
(A) Analysis of a Δ*fplA* mutant on *F. nucleatum* 23726 growth in rich CBHK media reveals no growth defect in the abscence of FplA. (B-D) Loss of FplA does not alter bacterial cell shape or length as deter mined by analyzing n=∼500 bacteria using the MicrobeJ plugin for lmageJ. Statistical analysis was performed using an unpaired t test. ns = not significant (p-value = 0.195).

**Fig S6.**
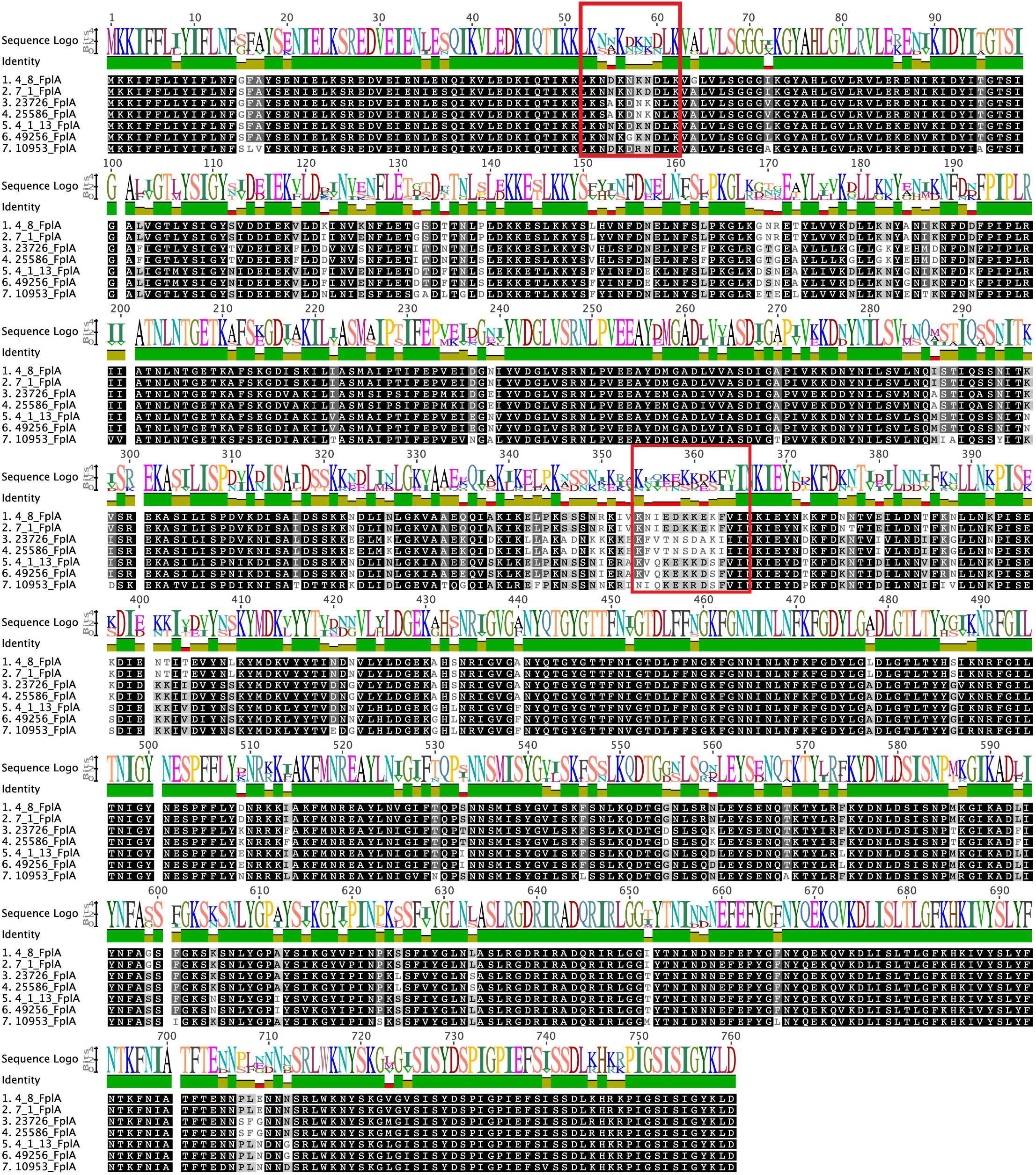
Alignment of full-length FplA from 7 strains of Fusobacterium and potential cleavage sites for release of the PLA, domain are outlined in red and were determined based on differences seen in the 23726 and 25586 strains (not cleaved) when compared to the remaining five strains that can be cleaved.

**Fig S7.**
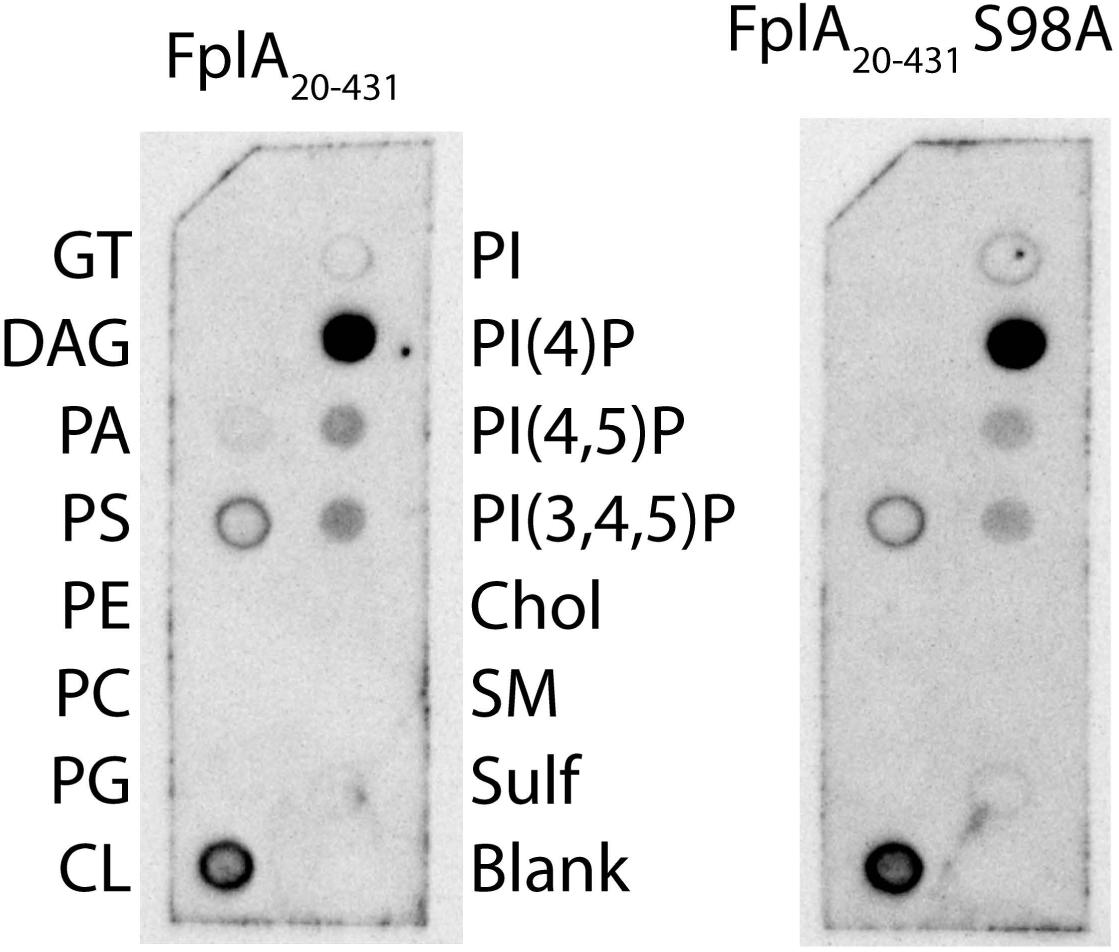
Initial analysis of FplA_20-431_ and FplA_20-431_ S98A lipid binding using commercially available lipid strips from Eschelon Inc. Subsequent analysis used freshly blotted lipids (Avanti Polar Lipids) and revealed strong affinity for phosphoinsositides, but not P1(4)p as seen here.

**supplementary Table 1:** Bacterial strains used in this study (Casasanta et al).

| Strain | Bacterial Species | Relevant genotype | Source or Reference |
| --- | --- | --- | --- |
| TOP10 | *E. coli* | *mcrA*, $\Delta(mrr-hsdRMS-mcrBC)$, Φ80(*del*)M15, ΔlacX74, *deoR*, *recA1*, *araD139*, $\Delta(ara-leu)$7697, *galU*, *galK*, *rpsL*(*SmR*), *endA1*, *nupG* | Invitrogen |
| LOBSTR-BL21(DE3)-RIL | *E. coli* | *fhuA2* [*lon*] *ompT gal* ($\lambda$ DE3) [*dcm*] Δ*hsdS* $\lambda$ DE3 = $\lambda$ *sBamHlo* Δ*EcoRI-B int::(lacI::PlacUV5::T7 gene1)* *i21* Δ*nin5* *arnA*(H359S, H361S, H592S, H593S) *slyD*(1-150) | [1] |
| *F. nucleatum nucleatum* ATCC 23726 | *F. nucleatum* | Wild Type | ATCC, [2–5] |
| DJSVT01 | *F. nucleatum* | *F. nucleatum* ATCC 23726 *Cm<sup>r</sup> Tm<sup>r</sup> ΔFN1704::catP* | This Study |
| *F. nucleatum nucleatum* ATCC 25586 | *F. nucleatum* | Wild Type | ATCC, [2–5] |
| *F. nucleatum polymorphum* 10953 | *F. nucleatum* | Wild Type | ATCC, [2–5] |
| *F. nucleatum vincentii* ATCC 49256 | *F. nucleatum* | Wild Type | ATCC, [2–5] |
| *F. nucleatum vincentii* 4_1_13 | *F. nucleatum* | Wild Type | [5] |
| *F. nucleatum animalis* 4_8 | *F. nucleatum* | Wild Type | [5] |
| *F. nucleatum animalis* 7_1 | *F. nucleatum* | Wild Type | [5] |

*Cm<sup>r</sup>*, Chloramphenicol resistance
*Tm<sup>r</sup>*, Thiamphenicol resistance
- Andersen KR, Leksa NC, Schwartz TU. Optimized *E. coli* expression strain LOBSTR eliminates common contaminants from His-tag purification. *Proteins*. 2013;81: 1857–1861. - Knorr M. Über die fusospirilläre Symbiose, die Gattung *Fusobacterium* (KB Lehmann) und *Spirillum sputigenum*. Zugleich ein Beitrag zur Bakteriologie der Mundhöhle. II. Mitteilung Die Gattung *Fusobacterium* I Abt Orig Zentralbl Bakteriол Parasitenkd Infektionskr Hyg. 1922;89: 4–22. - Knorr M. Ueber die fusospirilläre symbiose, die Gattung *Fusobacterium* (KB Lehmann) und *Spirillum sputigenum*. II Mitteilung. Die Gattung *Fusobacterium*. Zentbl Bakteriол Parasitenkd Infekt Hyg Abt. 1923;1: 4–22. - Dzink JL, Sheenan MT, Socransky SS. Proposal of three subspecies of *Fusobacterium nucleatum* Knorr 1922: *Fusobacterium nucleatum* subsp. *nucleatum* subsp. nov., comb. nov.; *Fusobacterium nucleatum* subsp. *polymorphum* subsp. nov., nom. rev., comb. nov.; and *Fusobacterium nucleatum* subsp. *vincentii* subsp. nov., nom. rev., comb. nov. *Int J Syst Evol Microbiol*. Microbiology Society; 1990;40: 74–78. - Manson McGuire A, Cochrane K, Griggs AD, Haas BJ, Abeel T, Zeng Q, et al. Evolution of invasion in a diverse set of *Fusobacterium* species. *MBio*. 2014;5: e01864.

**supplementary Table 2:** Plasmids used in this study (Casasanta et al).

| Plasmid | Description | Reference |
| --- | --- | --- |
| pET16b | *E. coli* inducible expression vector | EMD Millipore |
| pJIR750 | Base *C. perfringens*-*E. coli* shuttle vector to make *F. nucleatum* shuttle vectors. | [1] |
| pDJSVT43 | *F. nucleatum* 25586 FN1704 (*fplA*) 20-431 C-6xHis cloned into pET16b | This Study |
| pDJSVT60 | *F. nucleatum* 25586 FN1704 (*fplA*) 20-431 S98A C-6xHis cloned into pET16b | This Study |
| pDJSVT61 | *F. nucleatum* 25586 FN1704 (*fplA*) 20-431 D243A C-6xHis cloned into pET16b | This Study |
| pDJSVT82 | *F. nucleatum* 25586 FN1704 (*fplA*) 60-431 C-6xHis cloned into pET16b | This Study |
| pDJSVT84 | *F. nucleatum* 25586 FN1704 (*fplA*) 20-350 C-6xHis cloned into pET16b | This Study |
| pDJSVT85 | *F. nucleatum* 25586 FN1704 (*fplA*) 60-350 C-6xHis cloned into pET16b | This Study |
| pDJSVT86 | *E. coli* inducible expression vector - pET16b with *E. coli* OmpA signal sequence (residues 1-27) followed by a 6xHis tag and multiple cloning site. | This Study |
| pDJSVT88 | *F. nucleatum* 25586 FN1704 (*fplA*) 20-760 cloned into pDJSVT86 for expression of full length FplA on the surface of *E. coli* | This Study |
| pDJSVT100 | Shuttle vector to make strain DJSVT01 (*Table S1*): *F. nucleatum* ATCC 23726 ΔFN1704::*catP* (*Cm<sup>r</sup>* *Tm<sup>r</sup>*) | This Study |

*Cm<sup>r</sup>*, Chloramphenicol resistance
*Tm<sup>r</sup>*, Thiamphenicol resistance
1. Bannam TL, Rood JI. Clostridium perfringens-Escherichia coli shuttle vectors that carry single antibiotic resistance determinants. Plasmid. 1993;29: 233–235.

**supplementary Table 3:**
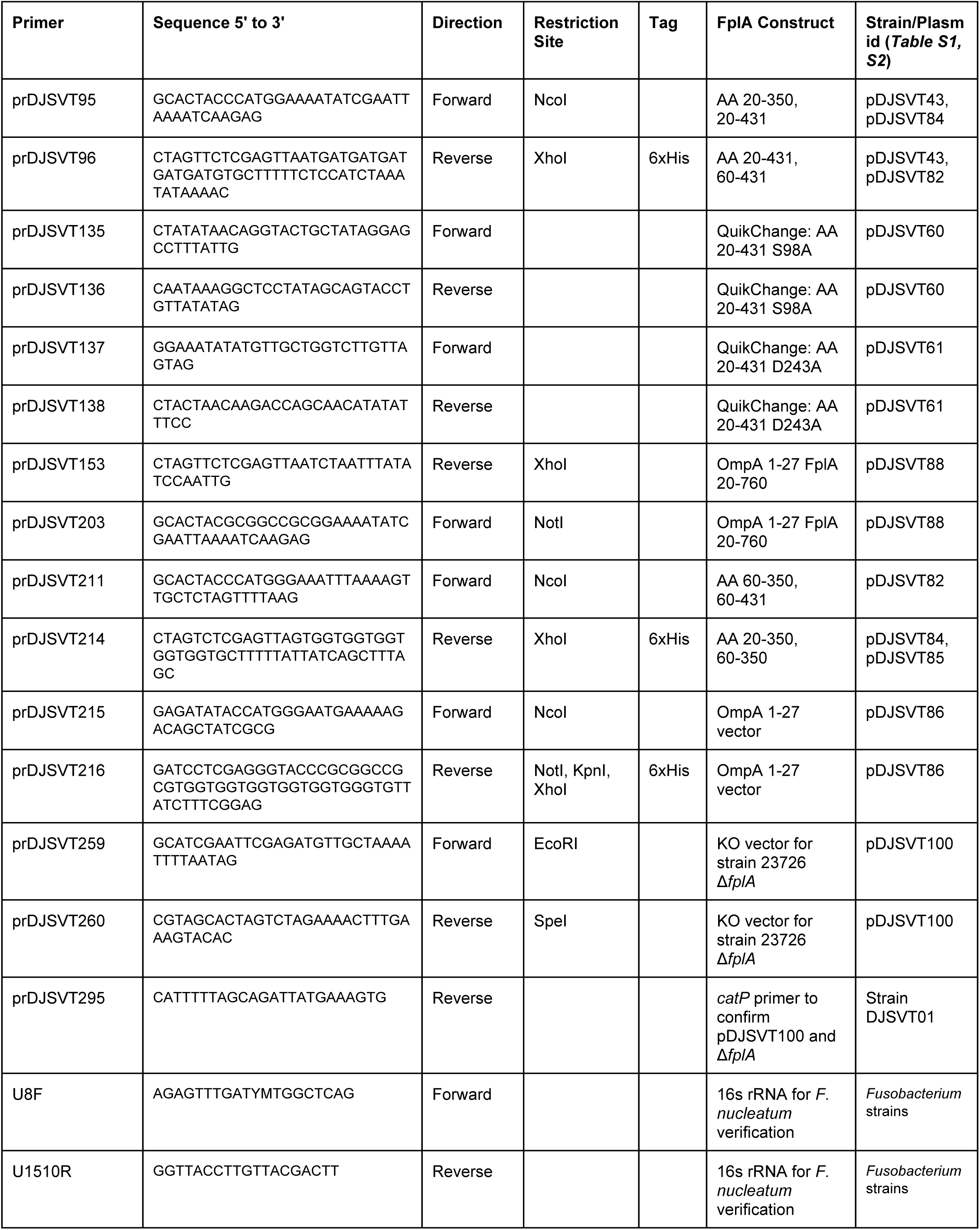
Primers introduced in this study (Casasanta et al).

## REFERENCES

1. Han, Y. W. et al. *Fusobacterium nucleatum* induces premature and term stillbirths in pregnant mice: implication of oral bacteria in preterm birth. Infect. Immun. 72, 2272–2279 (2004).

2. Abed, J. et al. Fap2 Mediates *Fusobacterium nucleatum* Colorectal Adenocarcinoma Enrichment by Binding to Tumor-Expressed Gal-GalNAc. Cell Host Microbe 20, 215–225 (2016).

3. Dahya, V., Patel, J., Wheeler, M. & Ketsela, G. Fusobacterium nucleatum endocarditis presenting as liver and brain abscesses in an immunocompetent patient. Am. J. Med. Sci. 349, 284–285 (2015).

4. Rashidi, A., Tahhan, S. G., Cohee, M. W. & Goodman, B. M. *Fusobacterium nucleatum* infection mimicking metastatic cancer. Indian J. Gastroenterol. 31, 198–200 (2012).

5. Gedik, A. H., Cakir, E., Soysal, O. & Umutoğlu, T. Endobronchial lesion due to pulmonary *Fusobacterium nucleatum* infection in a child. Pediatr. Pulmonol. 49, E63–5 (2014).

6. Shammas, N. W. et al. Infective endocarditis due to *Fusobacterium nucleatum*: case report and review of the literature. Clin. Cardiol. 16, 72–75 (1993).

7. Swidsinski, A. et al. Acute appendicitis is characterised by local invasion with *Fusobacterium nucleatum/necrophorum*. Gut 60, 34–40 (2011).

8. Han, Y. W. *Fusobacterium nucleatum*: a commensal-turned pathogen. Curr. Opin. Microbiol. 23, 141–147 (2015).

9. Gauthier, S. et al. The origin of *Fusobacterium nucleatum* involved in intra-amniotic infection and preterm birth. J. Matern. Fetal. Neonatal Med. 24, 1329–1332 (2011).

10. Castellarin, M. et al. *Fusobacterium nucleatum* infection is prevalent in human colorectal carcinoma. Genome Res. 22, 299–306 (2012).

11. Warren, R. L. et al. Co-occurrence of anaerobic bacteria in colorectal carcinomas. Microbiome 1, 16 (2013).

12. Kostic, A. D. et al. Genomic analysis identifies association of *Fusobacterium* with colorectal carcinoma. Genome Res. 22, 292–298 (2012).

13. Kostic, A. D. et al. *Fusobacterium nucleatum* potentiates intestinal tumorigenesis and modulates the tumor-immune microenvironment. Cell Host Microbe 14, 207–215 (2013).

14. Flanagan, L. et al. *Fusobacterium nucleatum* associates with stages of colorectal neoplasia development, colorectal cancer and disease outcome. Eur. J. Clin. Microbiol. Infect. Dis. 33, 1381–1390 (2014).

15. Strauss, J. et al. Invasive potential of gut mucosa-derived *Fusobacterium nucleatum* positively correlates with IBD status of the host. Inflamm. Bowel Dis. 17, 1971–1978 (2011).

16. Manson McGuire, A. et al. Evolution of invasion in a diverse set of *Fusobacterium* species. MBio 5, e01864 (2014).

17. Rubinstein, M. R. et al. *Fusobacterium nucleatum* promotes colorectal carcinogenesis by modulating E-cadherin/β-catenin signaling via its FadA adhesin. Cell Host Microbe 14, 195–206 (2013).

18. Xu, M. et al. FadA from *Fusobacterium nucleatum* utilizes both secreted and nonsecreted forms for functional oligomerization for attachment and invasion of host cells. J. Biol. Chem. 282, 25000–25009 (2007).

19. Kaplan, C. W. et al. *Fusobacterium nucleatum* outer membrane proteins Fap2 and RadD induce cell death in human lymphocytes. Infect. Immun. 78, 4773–4778 (2010).

20. Gur, C. et al. Binding of the Fap2 protein of *Fusobacterium nucleatum* to human inhibitory receptor TIGIT protects tumors from immune cell attack. Immunity 42, 344–355 (2015).

21. Coppenhagen-Glazer, S. et al. Fap2 of *Fusobacterium nucleatum* is a galactose-inhibitable adhesin involved in coaggregation, cell adhesion, and preterm birth. Infect. Immun. 83, 1104–1113 (2015).

22. Kaplan, C. W., Lux, R., Haake, S. K. & Shi, W. The Fusobacterium nucleatum outer membrane protein RadD is an arginine-inhibitable adhesin required for inter-species adherence and the structured. … Mol. Microbiol. (2009).

23. Gupta, S. et al. Fusobacterium nucleatum-associated beta-defensin inducer (FAD-I): identification, isolation, and functional evaluation. J. Biol. Chem. 285, 36523–36531 (2010).

24. Bhattacharyya, S. et al. FAD-I, a *Fusobacterium nucleatum* Cell Wall-Associated Diacylated Lipoprotein That Mediates Human Beta Defensin 2 Induction through Toll-Like Receptor-1/2 (TLR-1/2) and TLR-2/6. Infect. Immun. 84, 1446–1456 (2016).

25. Desvaux, M., Khan, A., Beatson, S. A., Scott-Tucker, A. & Henderson, I. R. Protein secretion systems in *Fusobacterium nucleatum*: genomic identification of Type 4 piliation and complete Type V pathways brings new insight into mechanisms of pathogenesis. Biochim. Biophys. Acta 1713, 92–112 (2005).

26. Henderson, I. R. & Nataro, J. P. Virulence functions of autotransporter proteins. Infect. Immun. 69, 1231–1243 (2001).

27. Wells, T. J., Tree, J. J., Ulett, G. C. & Schembri, M. A. Autotransporter proteins: novel targets at the bacterial cell surface. FEMS Microbiol. Lett. 274, 163–172 (2007).

28. Salacha, R. et al. The *Pseudomonas aeruginosa* patatin-like protein PlpD is the archetype of a novel Type V secretion system. Environ. Microbiol. 12, 1498–1512 (2010).

29. da Mata Madeira, P. V. et al. Structural Basis of Lipid Targeting and Destruction by the Type V Secretion System of *Pseudomonas aeruginosa*. J. Mol. Biol. 428, 1790–1803 (2016).

30. Pizarro-Cerdá, J. & Cossart, P. Subversion of phosphoinositide metabolism by intracellular bacterial pathogens. Nat. Cell Biol. 6, 1026–1033 (2004).

31. Istivan, T. S. & Coloe, P. J. Phospholipase A in Gram-negative bacteria and its role in pathogenesis. Microbiology 152, 1263–1274 (2006).

32. Flores-Díaz, M., Monturiol-Gross, L., Naylor, C., Alape-Girón, A. & Flieger, A. Bacterial Sphingomyelinases and Phospholipases as Virulence Factors. Microbiol. Mol. Biol. Rev. 80, 597–628 (2016).

33. Engel, J. & Balachandran, P. Role of *Pseudomonas aeruginosa* type III effectors in disease. Curr. Opin. Microbiol. 12, 61–66 (2009).

34. Tannaes, T., Bukholm, I. K. & Bukholm, G. High relative content of lysophospholipids of *Helicobacter pylori* mediates increased risk for ulcer disease. FEMS Immunol. Med. Microbiol. 44, 17–23 (2005).

35. Dorrell, N. et al. Characterization of *Helicobacter pylori* PldA, a phospholipase with a role in colonization of the gastric mucosa. Gastroenterology 117, 1098–1104 (1999).

36. Birmingham, C. L. et al. *Listeria monocytogenes* evades killing by autophagy during colonization of host cells. Autophagy 3, 442–451 (2007).

37. Anderson, D. M. et al. Ubiquitin and ubiquitin-modified proteins activate the *Pseudomonas aeruginosa* T3SS cytotoxin, ExoU. Mol. Microbiol. 82, 1454–1467 (2011).

38. Sato, H. & Frank, D. W. Intoxication of host cells by the T3SS phospholipase ExoU: PI(4,5)P2-associated, cytoskeletal collapse and late phase membrane blebbing. PLoS One 9, e103127 (2014).

39. Saliba, A. M. et al. Eicosanoid-mediated proinflammatory activity of *Pseudomonas aeruginosa* ExoU. Cell. Microbiol. 7, 1811–1822 (2005).

40. Lucas, M. et al. Structural basis for the recruitment and activation of the *Legionella* phospholipase VipD by the host GTPase Rab5. Proc. Natl. Acad. Sci. U. S. A. 111, E3514– 23 (2014).

41. Gaspar, A. H. & Machner, M. P. VipD is a Rab5-activated phospholipase A1 that protects *Legionella pneumophila* from endosomal fusion. Proc. Natl. Acad. Sci. U. S. A. 111, 4560– 4565 (2014).

42. Zhu, W., Hammad, L. A., Hsu, F., Mao, Y. & Luo, Z.-Q. Induction of caspase 3 activation by multiple Legionella pneumophila Dot/Icm substrates. Cell. Microbiol. 15, 1783–1795 (2013).

43. Biasini, M. et al. SWISS-MODEL: modelling protein tertiary and quaternary structure using evolutionary information. Nucleic Acids Res. 42, W252–8 (2014).

44. Lio, Y. C., Reynolds, L. J., Balsinde, J. & Dennis, E. A. Irreversible inhibition of Ca(2+)-independent phospholipase A2 by methyl arachidonyl fluorophosphonate. Biochim. Biophys. Acta 1302, 55–60 (1996).

45. Schevitz, R. W. et al. Structure-based design of the first potent and selective inhibitor of human non-pancreatic secretory phospholipase A2. Nat. Struct. Biol. 2, 458–465 (1995).

46. Lombardo, D. & Dennis, E. A. Cobra venom phospholipase A2 inhibition by manoalide. A novel type of phospholipase inhibitor. J. Biol. Chem. 260, 7234–7240 (1985).

47. Hope, W. C., Chen, T. & Morgan, D. W. Secretory phospholipase A 2 inhibitors and calmodulin antagonists as inhibitors of cytosolic phospholipase A 2. Inflamm. Res. 39, C39–C42 (1993).

48. Bennett, C. F. et al. Inhibition of phosphoinositide-specific phospholipase C by manoalide. Mol. Pharmacol. 32, 587–593 (1987).

49. Liu, Y., Patricelli, M. P. & Cravatt, B. F. Activity-based protein profiling: the serine hydrolases. Proc. Natl. Acad. Sci. U. S. A. 96, 14694–14699 (1999).

50. Simon, G. M. & Cravatt, B. F. Activity-based proteomics of enzyme superfamilies: serine hydrolases as a case study. J. Biol. Chem. 285, 11051–11055 (2010).

51. Dekker, N., Tommassen, J., Lustig, A., Rosenbusch, J. P. & Verheij, H. M. Dimerization regulates the enzymatic activity of *Escherichia coli* outer membrane phospholipase A. J. Biol. Chem. 272, 3179–3184 (1997).

52. Kinder Haake, S., Yoder, S. & Gerardo, S. H. Efficient gene transfer and targeted mutagenesis in *Fusobacterium nucleatum*. Plasmid 55, 27–38 (2006).

53. Bannam, T. L. & Rood, J. I. *Clostridium perfringens-Escherichia coli* shuttle vectors that carry single antibiotic resistance determinants. Plasmid 29, 233–235 (1993).

54. Sato, H. & Frank, D. W. ExoU is a potent intracellular phospholipase. Mol. Microbiol. 53, 1279–1290 (2004).

55. Howell, H. A., Logan, L. K. & Hauser, A. R. Type III secretion of ExoU is critical during early *Pseudomonas aeruginosa* pneumonia. MBio 4, e00032–13 (2013).

56. Machado, G.-B. S. et al. ExoU-induced vascular hyperpermeability and platelet activation in the course of experimental *Pseudomonas aeruginosa* pneumosepsis. Shock 33, 315–321 (2010).

57. Allewelt, M., Coleman, F. T., Grout, M., Priebe, G. P. & Pier, G. B. Acquisition of expression of the *Pseudomonas aeruginosa* ExoU cytotoxin leads to increased bacterial virulence in a murine model of acute pneumonia and systemic spread. Infect. Immun. 68, 3998–4004 (2000).

58. Afra, K., Laupland, K., Leal, J., Lloyd, T. & Gregson, D. Incidence, risk factors, and outcomes of *Fusobacterium* species bacteremia. BMC Infect. Dis. 13, 264 (2013).

59. Heckmann, J. G., Lang, C. J. G., Hartl, H. & Tomandl, B. Multiple brain abscesses caused by *Fusobacterium nucleatum* treated conservatively. Can. J. Neurol. Sci. 30, 266–268 (2003).

60. Ahmed, Z., Bansal, S. K. & Dhillon, S. Pyogenic liver abscess caused by *Fusobacterium* in a 21-year-old immunocompetent male. World J. Gastroenterol. 21, 3731–3735 (2015).

61. Brook, I. Infective endocarditis caused by anaerobic bacteria. Arch. Cardiovasc. Dis. 101, 665–676 (2008).

62. Agarwal, S. et al. Autophagy and endosomal trafficking inhibition by *Vibrio cholerae* MARTX toxin phosphatidylinositol-3-phosphate-specific phospholipase A1 activity. Nat. Commun. 6, 8745 (2015).

63. Gavin, H. E., Beubier, N. T. & Satchell, K. J. F. The Effector Domain Region of the Vibrio vulnificus MARTX Toxin Confers Biphasic Epithelial Barrier Disruption and Is Essential for Systemic Spread from the Intestine. PLoS Pathog. 13, e1006119 (2017).

64. Vergne, I. et al. Mechanism of phagolysosome biogenesis block by viable *Mycobacterium tuberculosis*. Proc. Natl. Acad. Sci. U. S. A. 102, 4033–4038 (2005).

65. Dong, N. et al. Modulation of membrane phosphoinositide dynamics by the phosphatidylinositide 4-kinase activity of the *Legionella* LepB effector. Nat Microbiol 2, 16236 (2016).

66. Muyzer, G., de Waal, E. C. & Uitterlinden, A. G. Profiling of complex microbial populations by denaturing gradient gel electrophoresis analysis of polymerase chain reaction-amplified genes coding for 16S rRNA. Appl. Environ. Microbiol. 59, 695–700 (1993).

67. Hyatt, D. et al. Prodigal: prokaryotic gene recognition and translation initiation site identification. BMC Bioinformatics 11, 119 (2010).

68. Eddy, S. R. Profile hidden Markov models. Bioinformatics 14, 755–763 (1998).

69. Kearse, M. et al. Geneious Basic: an integrated and extendable desktop software platform for the organization and analysis of sequence data. Bioinformatics 28, 1647–1649 (2012).

70. Arnold, K., Bordoli, L., Kopp, J. & Schwede, T. The SWISS-MODEL workspace: a web-based environment for protein structure homology modelling. Bioinformatics 22, 195–201 (2006).

71. Andersen, K. R., Leksa, N. C. & Schwartz, T. U. Optimized *E. coli* expression strain LOBSTR eliminates common contaminants from His-tag purification. Proteins 81, 1857– 1861 (2013).

72. Studier, F. W. Protein production by auto-induction in high density shaking cultures. Protein Expr. Purif. 41, 207–234 (2005).

73. Huang, Z., Liu, S., Street, I., Laliberte, F. & Abdullah, K. Methyl arachidonyl fluorophosphonate, a potent irreversible cPLA2 inhibitor, blocks the mobilization of arachidonic acid in human platelets and. … Mediators Inflamm. (1994).

74. Street, I. P. et al. Slow- and tight-binding inhibitors of the 85-kDa human phospholipase A2. Biochemistry 32, 5935–5940 (1993).

75. Segall, Y., Quistad, G. B., Sparks, S. E., Nomura, D. K. & Casida, J. E. Toxicological and structural features of organophosphorus and organosulfur cannabinoid CB1 receptor ligands. Toxicol. Sci. 76, 131–137 (2003).

76. Ackermann, E. J., Conde-Frieboes, K. & Dennis, E. A. Inhibition of Macrophage Ca-independent Phospholipase A by Bromoenol Lactone and Trifluoromethyl Ketones. J. Biol. Chem. 270, 445–450 (1995).

77. Adibekian, A., Martin, B. R., Speers, A. E., Brown, S. J. & Spicer, T. Optimization and characterization of a triazole urea dual inhibitor for lysophospholipase 1 (LYPLA1) and lysophospholipase 2 (LYPLA2). (2013).

78. Nomura, D. K. et al. Activation of the endocannabinoid system by organophosphorus nerve agents. Nat. Chem. Biol. 4, 373–378 (2008).

79. Randazzo, A. et al. Petrosaspongiolides M-R: new potent and selective phospholipase A2 inhibitors from the New Caledonian marine sponge *Petrosaspongia nigra*. J. Nat. Prod. 61, 571–575 (1998).

